# T-regulatory cell protection of progenitor cells from CD4^+^ T-cell-mediated cytotoxicity is essential for endogenous mouse digit-tip regeneration

**DOI:** 10.1101/2025.11.19.689270

**Authors:** Zachery Beal, Robyn E. Reeve, Elizabeth Hammond, Nadia Rosenthal, James Godwin

**Affiliations:** The Jackson Laboratory 600 Main Street Bar Harbor, ME 04609 USA; National Heart and Lung Institute Imperial Centre for Translational and Experimental Medicine Imperial College London, Faculty of Medicine Du Cane Road, London W12 0NN UK; MDI Biological Laboratory 159 Old Bar Harbor Rd Bar Harbor, ME 04609 USA

**Keywords:** regeneration, adaptive immunity, digit tip, T cell, microCT, CD4^+^, CD8^+^, T-reg, bone, FoxP3, osteoclasts, osteoblasts

## Abstract

Regeneration of amputated digit tips in humans and mice relies on osteoclast-dependent bone erosion coupled with osteoblast-mediated bone replacement. Currently, little is known of the impact of lymphoid immune cells, i.e., T cells, B cells, and NK cells, on digit-tip regeneration. Using lymphoid-deficient mutant mice, we revealed lymphoid immunity as a net negative regulator of regeneration. CD8^+^ cells are thought to negatively regulate fracture repair; however, we showed that adoptive cell transfer (ACT) of CD8^+^ T cells into lymphoid-deficient hosts did not impact regeneration. In contrast, ACT of CD4^+^ T cells potently inhibited regeneration via osteoclast and osteoblast progenitor-cell cytotoxicity. CD4^+^ T-cell-mediated inhibition of regeneration was rescued by supplementation with T regulatory cells or recombinant RANKL, a mediator of osteoclast differentiation. ACT of IFN-γ-deficient CD4^+^ T cells abolished cytotoxic activity and rescued regeneration. Future strategies protecting endogenous progenitor cells could enhance human tissue repair and autologous stem-cell therapies.

**One sentence summary:** Endogenous progenitor cells are vulnerable to CD4^+^ T-cell-mediated cytotoxicity during digit-tip regeneration and require T-regulatory-cell-mediated protection from autoimmune attack.

**Highlights:**

- Digit-tip regeneration is enhanced with the loss of lymphoid immunity.
- Regeneration requires T regulatory cells (T-regs) for maintenance of osteoclastogenesis when other lymphoid cells are present.
- T-regs enhance regeneration in the absence of lymphoid immunity during the anabolic phase.
- Like thymic NK cells, CD4^+^ T cells and not CD8^+^ T cells are responsible for inhibition of regeneration.
- RANKL is essential to the rate-limiting catabolic phase of digit-tip regeneration.
- Both T-regs and recombinant RANKL can rescue CD4^+^ T-cell inhibition.
- Genetic knockout of key cytotoxicity genes (IFNγ, Prf1, and TNFα) in immune-competent mice enhances regeneration.
- CD4^+^ T-cell ACT induces both apoptosis and necroptosis.
- CD4^+^ T-cell cytotoxicity is dependent on IFNγ.

## Introduction

The mouse digit tip represents a unique platform for studying endogenous adult tissue regeneration and the interactions among the cells required for regeneration, including circulating immune cells and endogenous stem cells. The ability to regenerate digit tips is conserved between mice and humans and is restricted to the distal 1/3 of the terminal phalanx (P3), with more proximal amputations resulting in failed regeneration^1^ ^2^ ^3^. Interestingly, regeneration in the mouse P3-digit regeneration model proceeds via intramembranous ossification that builds new bone without the cartilage intermediates and subsequent gradual replacement of cartilage with bone, i.e., endochondral ossification, used in fracture healing and embryonic digit development^4^. After P3-level amputation in mice, the wound-healing response recruits several immune-cell types that limit the risk of infection and promote wound closure. Monocytes, a type of innate myeloid immune cell, dominate the early response, during which they differentiate into macrophages (Mϕs) and osteoclasts (OCs), the latter of which are cells that degrade bone via resorption. Regeneration after P3-level amputations proceeds via OC-mediated degradation of the P3 bone between 6 and 12 days post amputation^5^. This process triggers the recruitment of stem cell-like progenitors, some of which undergo differentiation directly into osteoblast (OB) cells^4^, a cell type that builds new bone and remodels existing bone. Importantly, both bone development and bone regeneration require the coordinated, balanced functioning of OC cells and OB cells. The importance of myeloid cells in digit tip regeneration is reflected in work showing that Mϕ and OC depletion in mice inhibits P3-digit regeneration^6^. Furthermore, monocytes and Mϕs are critical for successful repair in many different tissue types and animal models^7^ ^8^ ^9^ ^10^ including mouse fracture healing^11^.

Lymphoid immune cells are regulators of bone homeostasis, disease and repair^12^ but have not been well studied in epimorphic digit tip regeneration, a type of repair that forms a mass of undifferentiated cells called a blastema, which is absent in fracture repair^5^. Innate lymphoid cells i.e., natural killer (NK) cells, develop without antigen specificity and with germline-encoded receptors distinguishing them from adaptive lymphoid cells (i.e., T and B cells) who generate unique non-germline antigen receptors via rearrangement of antigen-receptor genes, mediated by V(D)J recombination-activating genes (RAGs) during lymphocyte development^13^. Although NK cells trigger osteoclastogenesis and bone destruction in arthritis models^14^ little is known about the role of NK cells in fracture healing^15^. We previously discovered that different NK cell subsets play both positive and negative roles in endogenous P3 digit-tip regeneration in mice^16^. Specifically, we showed that an NK subpopulation that develops in the thymus actively kills both OC and OB progenitor cells when unconstrained, whereas spleen-derived NK cells play a protective role^16^. Conversely, adaptive lymphoid cells (i.e., T cells, and B cells) have emerged as critical regulators of endochondral fracture healing by controlling OB and OC function^12^ but their roles have not been defined in intramembranous digit tip regeneration.

Different T cell subsets can have opposing functions in fracture repair^12^. T cells develop via specific antigen signaling in the thymus where they differentiate into either CD4^+^ or CD8^+^ cells. Most CD8^+^ T cells are cytotoxic lymphocytes (CTLs) that express receptors that can recognize a specific antigen. Mature naïve CD4^+^ T cells are helper T cells that, upon antigen stimulation in the periphery, can transform into at least six subtype of effector cells with distinct functions based on their microenvironment^17^. T regulatory cells (T-regs) are a particularly important CD4^+^ subset, playing essential roles in maintaining self-tolerance and both bone and immune-system homeostasis. CD8^+^ T cells, which are cytotoxic to cancer cells in mice and humans^18^, are implicated in delaying fracture healing in humans and have been shown to directly inhibit fracture healing in mice^19^. Together, the above evidence suggests that T cells likely influence mouse digit-tip regeneration; however, the roles of the major T-cell subtypes are not yet defined.

To determine the requirement for lymphoid immunity in mouse digit-tip regeneration and define the roles of the different T-cell subtypes, we used 16 mouse strains, including lymphoid-deficient and genetically modified mice, to examine the roles of specific lymphoid cell types in regeneration. The 16 strains included mice on three distinct genetic backgrounds (C57BL/6J, NOD and NU/J), enabling us to assess whether genetic background impacts regeneration. Our major finding is that, unexpectedly, endogenous P3 digit-tip regeneration performed best in the absence of all lymphoid immune cells: Whereas CD8^+^ T cells did not inhibit regeneration, CD4^+^ T cells functioned as potent inhibitors of the regenerative response. Specifically, CD4^+^ T cells induced apoptosis and necroptosis of OC progenitors required in the initial rate-limiting catabolic step of bone regeneration and inhibited the OB progenitor induction needed for anabolic bone growth. In contrast, adoptive transfer of T-regs into lymphoid-cell-deficient hosts enhanced regeneration during the OB-driven anabolic phase and played a role in maintaining regeneration in the presence of normal immunity, in part by suppressing CD4^+^ T-cell-mediated cytotoxicity. We also showed that the cytotoxic genes *IFN-γ, Prf1, and TNF-α* are involved in CD4^+^ T-cell-mediated inhibition of regeneration, with *IFN-γ* playing a major role. Overall, our study demonstrates that lymphoid immune cells have the capacity to destroy progenitor cells required for endogenous regeneration, a finding with profound implications for regenerative medicine. Although the immune system is primarily for protecting the organism from infection and cancer, endogenous regeneration requires a successful balance between the protective and destructive components of the lymphoid immune system.

## Results

### T cells are recruited to the early regenerating P3 digit tip

Previous work shows that during the first six days post-amputation (DPA) at the P3 position (distal 1/3 of terminal phalanx) in the mouse digit tip, wound healing and epithelial closure take place without any significant bone catabolism^16^. To assess T-cell recruitment during this early wound-healing phase, we performed confocal microscopy on amputated digits of wildtype C57BL6 (B6) mice at 6 DPA and immunostained the CD3 component of the T-cell receptor expressed on all T cells **(Fig. 1A, B)** or measured the recruitment of adoptively transferred red fluorescent (tdTomato^+^) T cells in B6RG **(Fig. 1C)**. Both methods confirmed that T cells are robustly recruited to amputated digits by 6 DPA.

**Figure 1.**
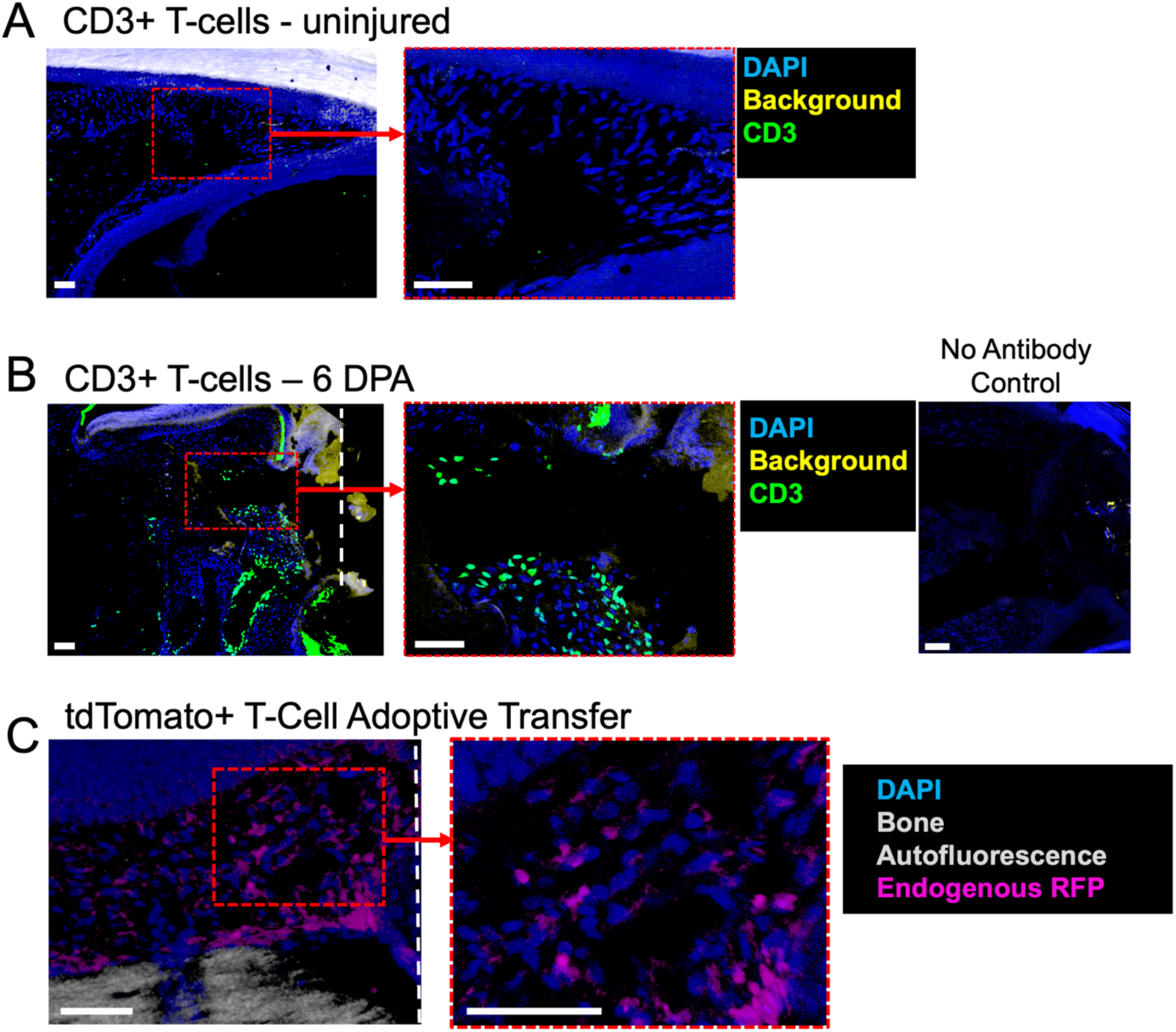
Mouse digit-tip amputation recruits endogenous T cells and adoptively transferred T cells into the injury site by 6 DPA. **A.** Low T-cell numbers in uninjured C57BL6/J (B6) mouse digit-tips. **B.** Robust T-cell recruitment to P3-level digit-tip amputations in B6 mice. **C.** Magnetically purified donor TdTomato^+^ CD3^+^ T cells migrate to the injured digit tip by 6 DPA after ACT at -1DPA in B6RG mice. A, B, and C: Red square in left panel shows inset. Approximate amputation plane is shown in white dashed line. Scale bar = 50 µm.

### Mice lacking thymic development of lymphoid cells show enhanced regeneration

In mice, adult digit-tip regeneration is distinct from embryonic digit-tip regeneration in that it undergoes a bone histolytic response followed by osteogenesis and new bone formation without the use of a cartilage intermediate^20^. To monitor and quantify the progress of both the catabolic histolytic step and the anabolic new bone formation, we used microcomputed tomography (microCT) to measure “hard-tissue” (ossified bone) volume, which usually completes by 12 DPA, and measured the increase in “soft-tissue” volume (which includes nail epithelium, and connective tissue mesenchyme) associated with formation of the blastema, a mass of highly proliferative cells^4^. As the blastema differentiates in the P3 mouse digit-tip model, the hard-tissue volume increases and the soft-tissue volume decreases. Our new thresholding approach for measuring hard and soft tissue in parallel allows improved visualization of the regenerating digit compared to previous microCT analysis methods^5^ introducing software that can quantify 3D volumes of hard or soft tissue or generate 3D renderings of bone in white and soft tissue in teal (**Fig. S1**).

To investigate the role of T cells in regulating the digit-tip regenerative response, we tested regrowth post-amputation in athymic nude mice (NU/J, spontaneous Foxn1*^nu^* mutation on a “nude-specific” inbred genetic background), and B6-nude mice (B6-Foxn1*^nu^*mice, where the *nu* mutation is on an inbred C57BL6/J background). Both of these “nude” mouse strains fail to develop the thymic organ, which is necessary for T-cell development, resulting in an absence of mature T cells, B cells, and the NK subpopulation that develops in the thymus. Despite the genetic background differences of the two athymic nude mouse strains, both exhibited early entry into the necessary bone-catabolism phase, followed by a near immediate transition to a rapid bone-outgrowth phase **(Fig. 2A, B, C).** Bone catabolism was complete by 6 DPA in both strains of nude mice, whereas this phase was not completed until 12 DPA in control B6 mice. In addition, both nude strains showed an early burst of soft-tissue outgrowth **(Fig. 2C)** associated with catabolism. To investigate the impact of T cells on these processes we took advantage of the back biopsy skin wound-healing model in hairless nude (NU/J) mice, which are T-cell-deficient, and the high cell yield attainable with this approach, which facilitates high-dimensional immunophenotyping. Adoptive cell transfer (ACT) of CD3^+^ T cells into NU/J mice resulted in a dramatic shift in myeloid-cell phenotypic diversity in skin wounds (lacking bone) **(Fig. S2A, B),** suggesting that T cells have a more general role in shaping the wound-healing response via myeloid-cell differentiation or polarization.

**Figure 2.**
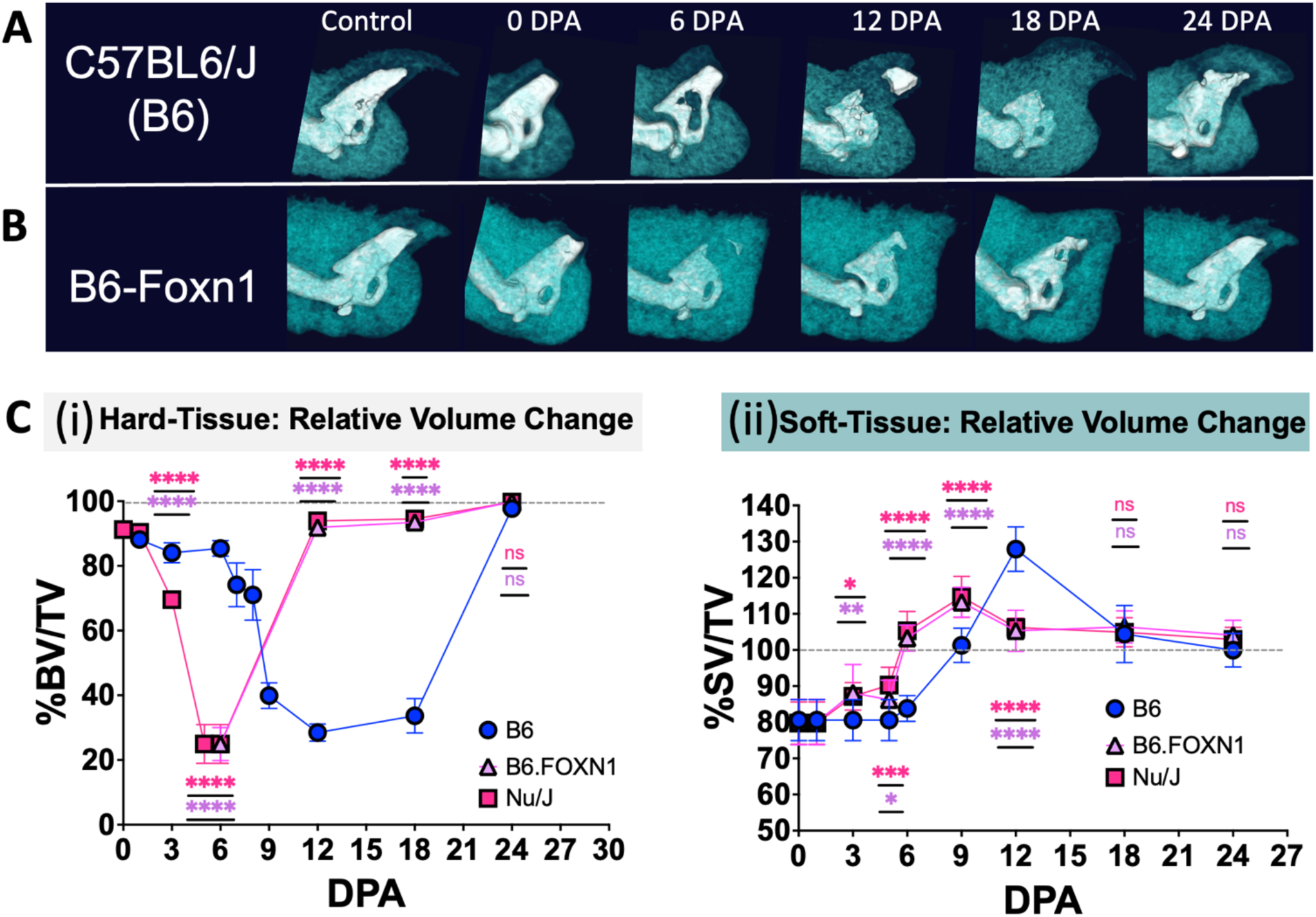
Both strains of athymic Foxn1^-/-^ (nude) mice, which lack T cells, B cells, and thymic NK cells, show accelerated regeneration, suggesting that, in digit-tip regeneration, lymphoid cells are dispensable and have a net inhibitory role. Digits of C57BL6/J (B6) mice, used as a control strain; B6-Foxn1 *^nu^* mice (nude on B6 genetic background); and NU/J mice (nude genetic background) were amputated at the P3 position and then assessed for hard- and soft-tissue regeneration over time, relative to unamputated control digits. **A, B**. Representative two-tone microCT images showing regeneration over time after P3-level amputations in B6 and B6-Foxn1 mice. Bone (hard tissue) shown in white, and soft tissue shown in teal. **C**. Both B6-Foxn1 and NU/J nude mice exhibit enhanced digit-tip regeneration with accelerated bone histolysis and new bone formation. Changes in hard tissue (i) and soft tissue (ii) relative to uninjured control over time. N = 5 mice with 4 amputated digits per animal = 20 measurements per condition, per timepoint. Data representative of at least 2 independent experiments. BV = bone volume, SV = soft volume, TV = total volume. P values calculated using one-way ANOVA with Tukey’s multiple comparison test. Statistical differences relative to B6 reference strain are indicated as ^ns^P > 0.05, *P ≤ 0.05, **P ≤ 0.01, ***P ≤ 0.001, ****P ≤ 0.0001. Means and standard deviations are shown.

### Digit-tip regeneration speed is positively correlated with the loss of lymphoid immunity

While nude mice are a classic model of mice lacking mature T cells, they also have defects in epithelial differentiation that may make unknown contributions to the phenotype^21^. To investigate the role of T cells in digit-tip regeneration in the absence of this potential confounding factor, we tested B6 mice harboring a Recombination Activating Gene-1 knockout (RAG-KO) mutation that results in a failure to produce mature T and B cells. Like nude mice, B6-RAG-KO mice have accelerated new bone formation in the anabolic phase and accelerated accumulation of soft-tissue outgrowth in the early wound-healing phase, but, unlike the nude strains, they do not show dramatically accelerated bone histolysis in the catabolic phase (**Fig. 3A, B**).

**Figure 3.**
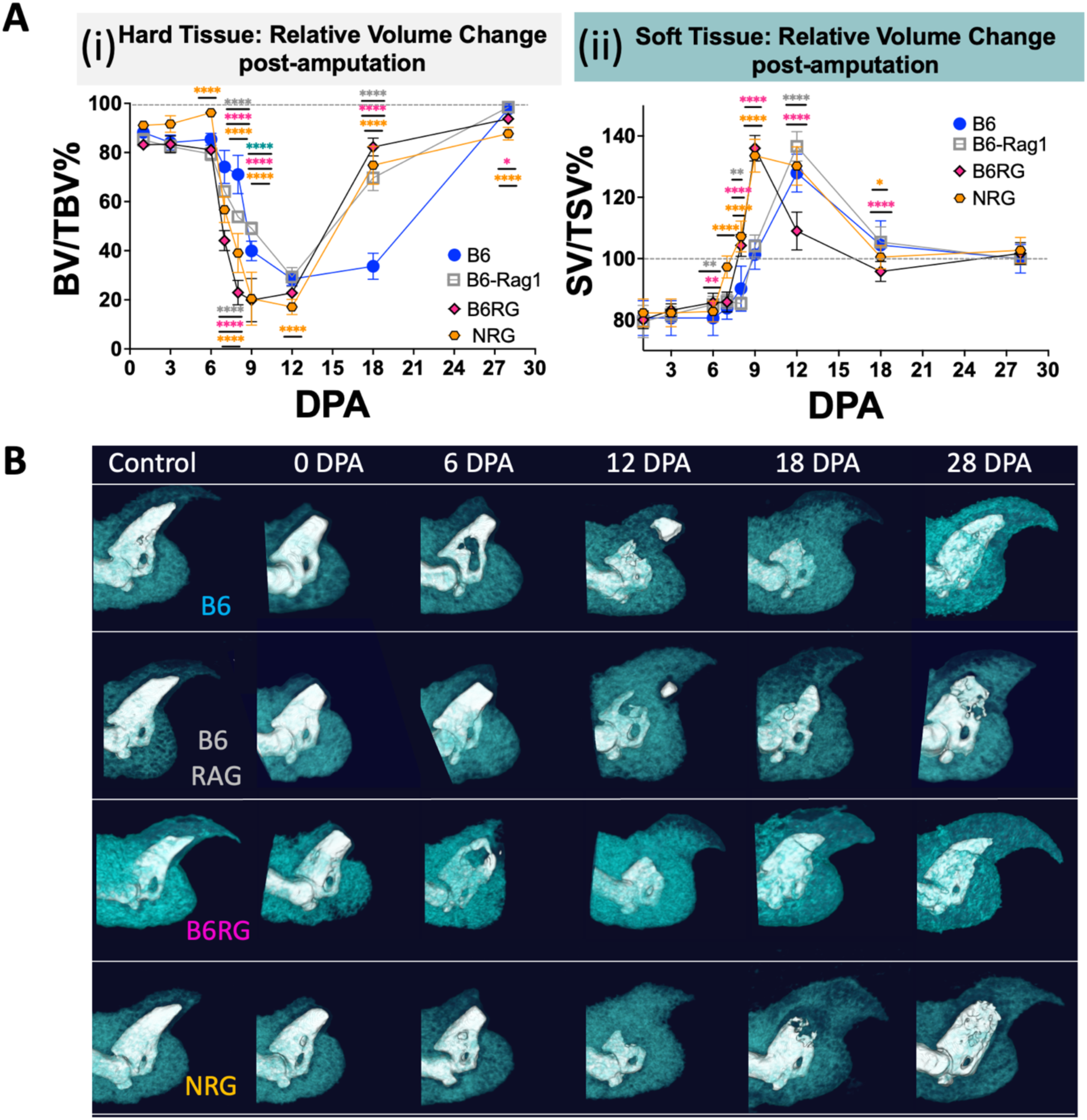
Mutant mice completely lacking lymphoid immune cells show enhanced regeneration in both catabolic and anabolic phases. Relative to C57BL6/J (B6) control mice, B6-RAG-1-KO mice, which lack T and B cells but retain NK cells, show enhanced anabolic bone growth in digit-tip regeneration. B6RG and NRG mice, which lack all lymphoid immune cells (T cells, B cells, NK cells), show enhancement in both the catabolic and anabolic phases of regeneration. **A.** Changes in hard tissue (i) and soft tissue (ii) relative to uninjured control over time. N = 5 mice, with 4 amputated digits per animal = 20 measurements per condition, per timepoint. Data representative of at least 2 independent experiments. BV = bone volume, SV = soft volume, TV = total volume. P values calculated using one-way ANOVA with Turkey’s multiple comparison test. Statistical differences relative to B6 reference strain indicated as ^ns^P > 0.05, *P ≤ 0.05, **P ≤ 0.01, ***P ≤ 0.001, ****P ≤ 0.0001. Means and standard deviations are shown. **B**. Representative two-tone microCT images of regeneration over time comparing strains. Bone (hard tissue) shown in white, and soft tissue shown in teal.

To determine whether NK cells impact bone catabolism, we crossed B6-RAG-2KO mice to interleukin 2 receptor gamma (IL2-Rγ) KO mice, which lack NK-cell development^22^. The strain resulting from this cross, which we termed “B6RG” mice, exhibited accelerated entry into and exit from the bone catabolic step compared to B6 or B6-RAG-1-KO mice, along with faster bone outgrowth the parental strains **(Fig. 3A, B).** To confirm these findings, we analyzed digit-tip regeneration in NRG mice, which completely lack adaptive/lymphoid immunity. NRG mice carry RAG-1 and IL2Rγ KO mutations that eliminate T-cell and NK-cell development on a non-obese diabetic genetic background (NOD) which confers additional immunological defects such as lower NK numbers, complement deficiencies and divergent macrophage responses^23^. Thus, both B6RG and NRG mice lack T, B, and NK cells, and while differing in their genetic backgrounds perform near identically in hard tissue regeneration with accelerated entry into the catabolic phase and rapid anabolic phase compared to B6 mice **(Fig. 3A, B)**. In addition, both B6RG and NRG mice showed an associated acceleration in soft-tissue outgrowth relative to B6 mice **(Fig. 3A, B)** confirming a net negative role for lymphoid cells in digit-tip regeneration.

### Effective regeneration requires T regulatory cells (T-regs) when other lymphoid cells are present

T-regs, defined by expression of the FoxP3 transcription factor, play important regulatory roles in immunity, regeneration, and bone homeostasis, promoting regeneration in several contexts through both cell contact and paracrine mechanisms (reviewed in^24, 25^). However, the molecular mechanisms underlying these functions are poorly understood, in part because of conflicting findings. For example, T-regs are known regulators of osteoclast formation in bone homeostasis and disease^26^, but studies suggest that they can both inhibit and promote osteoclastogenesis^27^ ^28^ ^29^ ^30^ ^31^. Osteoclast development is dependent upon stimulation of RANK^32^ (Receptor activator of NF-κB) on the surface of monocytes by its ligand RANKL. In many contexts, T-regs inhibit osteoclastogenesis via cytokine secretion^27^ ^28^, whereas in microenvironments rich in the pro-inflammatory cytokines IL-1β or IL-6 in mice or humans, T-regs can promote osteoclastogenesis via expression of RANKL^29^ ^30^ ^31^. To clarify the role of T-regs in the regenerating digit tip, we used FoxP3:DTR mice, which allows for selective systemic ablation of host T-regs by selectively expressing the Diphtheria Toxin (DT) receptor in T-regs. Injection of DT ablates T-regs and leaves non-T-reg cells undamaged in these mice. T-reg ablation over the time course of regeneration dramatically delayed initiation of the bone catabolic phase, from 6 DPA to 12 DPA, shifting the muted catabolic low to 18dpa. This also delayed the linked bone anabolic phase, with T-reg-depleted animals showing incomplete bone regeneration at 24 DPA, compared to the fully regenerated bone seen in control mice (vehicle-injected) at that time point (**Fig. 4A**). T-reg-depleted animals also showed severely impaired soft-tissue growth compared to control animals **(Fig. 4A)**, suggesting that T-regs may also regulate blastemal growth. Histological assessment of digit regeneration in T-reg-depleted FoxP3:DTR mice showed suppression of CD68^+^ multinucleated OC formation **(Fig. 4B-E)** and of phospho-histone H3 positive (PHH3^+^) mitotic (M-phase) cell numbers, despite robust entry into a proliferative state and cell cycle, as indicated by PCNA staining **(Fig. 4F-I)**. Together, these results suggest a supporting role for T-regs in digit-tip osteoclastogenesis and either progenitor-cell survival or completion of cell division during proliferation.

**Figure 4.**
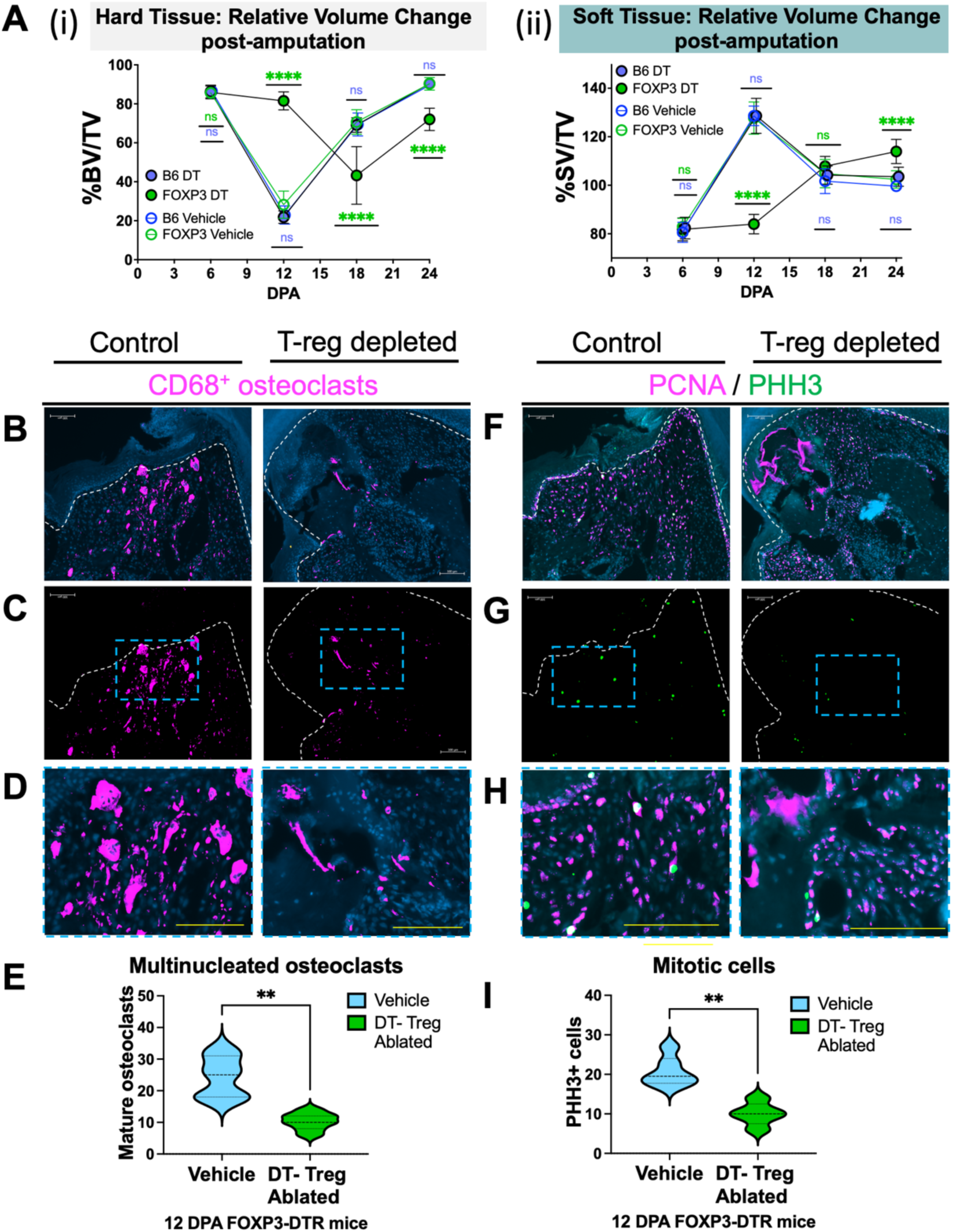
Diphtheria toxin (DT)-mediated depletion of genetically sensitized FoxP3^+^ T regulatory cells (T-regs) delays the bone-catabolism phase of regeneration and blastemal proliferation. **A**. Sustained DT-mediated ablation of T-regs in FoxP3-DTR mice results in disrupted digit-tip regeneration with delayed bone catabolism (i) and delayed soft-tissue growth (ii) (blastemal formation) relative to both vehicle-treated FoxP3-DTR mice or genetically insensitive (B6) controls. N = 5 mice, with 4 amputated digits per animal = 20 measurements per condition, per timepoint. Data representative of at least 3 independent experiments. BV = bone volume, SV = soft volume, TV = total volume. P values calculated using one-way ANOVA with Turkey’s multiple comparison test. **B-D.** Reduced numbers of mature multinucleated CD68^+^ osteoclasts observed in T-reg-depleted FoxP3:DTR mice (osteoclasts shown in magenta, nuclei in blue). **C.** Osteoclast channel alone with location of inset for D shown in blue dashed box. **E.** Quantification of mature osteoclast numbers. B, C. Boundary between blastema and nail epithelium shown in white dashed line. **F-H.** Reduced numbers of phospho-histone H3 (PHH3^+^) mitotic cells in T-reg-depleted FoxP3:DTR mice despite abundant PCNA proliferation marker expression (PCNA^+^ cells shown in magenta, PHH3^+^ in green, and nuclei in blue). **G.** PHH3^+^ channel alone with location of inset for **H** shown in blue dashed box. **I.** Quantification of PHH3^+^ cells in each condition. F, G. Boundary between blastema and nail epithelium shown in white dashed line. Statistical analysis via one-way ANOVA with Tukey post hoc. Differences relative to control are indicated as ^ns^P > 0.05, *P ≤ 0.05, **P ≤ 0.01, ***P ≤ 0.001, ****P ≤ 0.0001. Scale bar = 100 μm

Notably, systemic ablation of T-regs induced bone catabolism in uninjured control digits, both in the uninjured control digit 3 on both rear paws (with digits 2 and 4 amputated) and in the corresponding uninjured control digits on the front paws–**(Fig. S3A, B)**. These changes in hard-tissue volume were associated with OC induction, as indicated by the Tartrate-resistant acid phosphatase (TRAP^+^) marker^33^ (**Fig. S4A, B)**. Soft-tissue volume in uninjured control digits on both the rear and front paws was also increased in T-reg-deficient animals **(Fig. S3)** suggestive of hard-tissue break down or structural weakening and replacement with soft tissue. Taken together, these results show that T-reg ablation results in impaired osteoclastogenesis in the amputated digit-tip regeneration model, and in the induction of autoimmunity that promotes erosion of uninjured bone when other T cell subsets are present.

### T regulatory cells enhance regeneration in the absence of lymphoid immunity

To define the role of T-regs in the absence of other lymphoid cells, we first used ACT to transfer T-regs into the two lymphoid-deficient mouse strains used above, i.e., B6RG mice and NRG mice with different genetic backgrounds. For ACT experiments, we isolated T-regs from the mouse spleen, where approximately 10% of CD4^+^ T cells are CD4^+^FoxP3^+^ T-regs^34^. Using the FoxP3:YFP^+^ reporter mouse, we isolated splenic T-regs to 100% purity using FACS sorting and performed ACT into each strain prior to P3 digit amputation Live microCT was used to follow and quantify P3 digit-tip regeneration over time. Both strains exhibited T-reg-dependent acceleration of the OB-dependent anabolic bone-growth phase of regeneration, but no detectable impact of T-regs on the OC-dependent bone catabolic phase (**Fig. S5A**). T-reg-dependent anabolic acceleration is also associated with earlier soft tissue differentiation (**Fig. S5B**). However, while this ACT model simplifies the testing of target cells within a landscape lacking other adaptive lymphoid cells that could cloud interpretation, it may also lack other regulators that function within a normal immune repertoire, a scenario that must be considered when interpreting results.

### CD4^+^ T cells but not CD8^+^ T cells are potent inhibitors of regeneration

The results above confirm that adoptively transferred T cells can efficiently home to locations in the amputated mouse digit where endogenous T-cell invasion is also found **(Fig. 1)**. To directly test the relative contribution of CD4^+^ and CD8^+^ T cells in digit-tip regeneration, we used immunodeficient B6RG and NRG mice as host graft-accepting strains for ACT experiments **(Fig. S7B)**. We isolated CD4^+^ or CD8^+^ T cells from mouse spleens via magnetic negative selection, leaving these T cells “untouched” (i.e., no bound antibody) at over 96-98% purity and most likely unstimulated (**Fig. S6**)(avoiding positive selection that can sometimes trigger cellular changes and apoptosis). In both the B6RG **(Fig. 5A, B)** and NRG **(Fig. S7A, C, D)** genetic backgrounds, delivery of CD4^+^ T cells one day prior to amputation resulted in a substantial delay in bone catabolism; in the transition to the anabolic phase of new bone formation; and in soft-tissue outgrowth, that together caused major defects in regeneration. In contrast to the negative impact of both CD4^+^ T cells on digit-tip regeneration, ACT of CD8^+^ T cells into B6RG **(Fig. 5A, B)** and NRG (**Fig. S7A, C, D)** mice had no major effect on digit bone regeneration, which is surprising given that CD8^+^ cells are known to negatively regulate fracture repair^19^.

**Figure 5.**
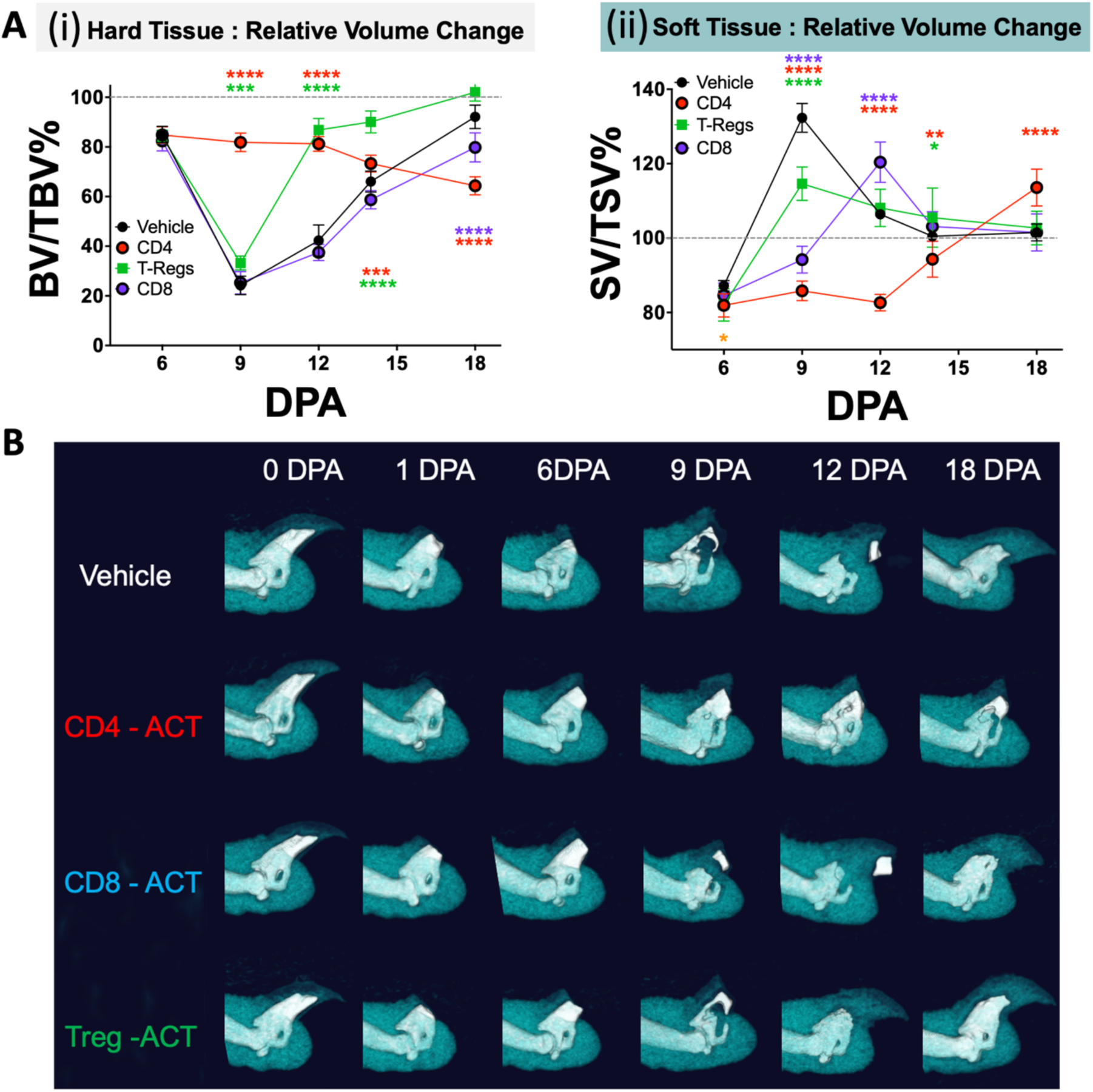
In B6RG mice lacking lymphoid immunity, ACT of naïve CD4^+^ T cells, but not of CD8^+^ T cells, blocks digit-tip regeneration, affecting both hard- and soft-tissue growth, whereas T-reg ACT enhances anabolic bone growth. The process of regeneration following ACT of magnetically purified CD4^+^ or CD8^+^ T cells or FACS-purified FoxP3^+/YFP+^ T-regs into B6RG mice one day prior to P3-level amputation differs depending on the cell type transferred. **A.** Changes in hard-tissue (i) and soft tissue (ii) relative to uninjured control digits over time. N = 5 mice with 4 amputated digits per animal = 20 measurements per condition, per timepoint. Data representative of at least 2 independent experiments. BV = bone volume, SV = soft volume, TV = total volume. P values calculated using one-way ANOVA with Turkey’s multiple comparison test. Statistical differences indicated as ^ns^P > 0.05, *P ≤ 0.05, **P ≤ 0.01, ***P ≤ 0.001, ****P ≤ 0.0001. Means and standard deviations are shown. Representative two-tone microCT images of regeneration over time comparing treatments. Bone (hard tissue) shown in white, and soft tissue shown in teal.

The defective regeneration caused by CD4+ T cell ACT is similar to the inhibition of digit-tip regeneration we reported previously using ACT of thymic NK (ThNK) cells into NSG mice^16^. To test the level of ThNK-cell-mediated inhibition of regeneration in the B6RG strain, we used homozygous Ncr1:GFP knock-in/knock-out mice expressing GFP exclusively in NK cells (lacking endogenous Ncr1 [NKp46]) to isolate GFP^+^ NK cells from both thymus and spleen with 100% purity using FACS sorting. ACT of ThNK cells showed potent inhibition of regeneration in B6RG comparable with the inhibition observed with CD4^+^ T cells **(Fig. 5, S8)**. These results aligned with our previous results in NSG mice^16^ and confirmed that *Ncr1* expression is essential for the pro-regenerative effects of splenic derived (SpNK) cells in B6RG mice.

### RANK ligand (RANKL) is essential to the rate-limiting osteoclast-dependent catabolic phase of digit-tip regeneration

To more precisely define the roles of lymphoid immune cells during the different phases of digit-tip regeneration, we tested whether bone histolysis is required for soft-tissue outgrowth and blastema formation by inhibiting RANKL function. As noted above, RANK and its ligand, RANKL, are key regulators of OC differentiation^32^*. In vivo* antibody neutralization of RANKL inhibits OC formation without affecting inflammation^35^. Neutralization of RANKL between 6 DPA and 18 DPA in wildtype B6 mice completely inhibited the catalytic histolytic step **(Fig. S9A, C, D)**; eliminated OC formation, as shown via quantification and visualization of TRAP^+^ osteoclasts **(Fig. S9B, C, respectively)**; and inhibited bone outgrowth, with no recovery by 40 DPA **(Fig. S9A, C, D).** RANKL neutralization also completely inhibited soft-tissue outgrowth **(Fig. S9A, C, D)**, indicating that histolysis of the amputated bone is critical for enabling soft-tissue outgrowth and blastema formation.

### Blockade of regeneration by CD4^+^ T-cell inhibition can be rescued by asynchronous T-reg or RANKL supplementation

Given the role of FoxP3^+^ T-regs in the regulation of other T-cell subsets and their important role in maintaining osteoclastogenesis (**Fig. 4)**, we explored the relationship between CD4^+^ T-cell-mediated inhibition of regeneration, and pro-regenerative T-reg activity. Although T-regs have been associated with suppression of OC production in some contexts^26, 36, 37^, they are also potent regulators of other CD4^+^ T-cell functions including Th1 and Th2 cytokine production and survival^38, 39, 40^. To test if T-regs could antagonize the inhibitory function of naïve CD4^+^ T cells, we performed ACT of purified CD4^+^ T cells one day prior to P3-level digit amputation in lymphoid-deficient B6RG and NRG mice, followed by an ACT chase with purified FoxP3^+^ T-regs 5DPA. Injection of T-regs at 5 DPA (6 days after CD4^+^ T-cell adoptive transfer) completely reversed the inhibitory role of CD4^+^ T cells in both the B6RG and NRG genetic backgrounds, rescuing regeneration entirely and bringing the catabolic and anabolic phases in line with control vehicle-injected mice **(Fig. 6A, B; Fig. S10A)**. We then tested whether the CD4^+^-mediated inhibition of regeneration is upstream of osteoclastogenesis, by determining if this inhibition is directly related to RANKL-dependent osteoclastogenesis. By delivering recombinant RANKL at 6 DPA, 8 DPA, and 10 DPA, we completely reversed the CD4^+^ T-cell-mediated inhibition of OC formation, bringing OC numbers back within normal range **(Fig. S10B, C)** and restoring full regeneration in lymphoid-deficient mice **(Fig. S10A, B).**

**Figure 6.**
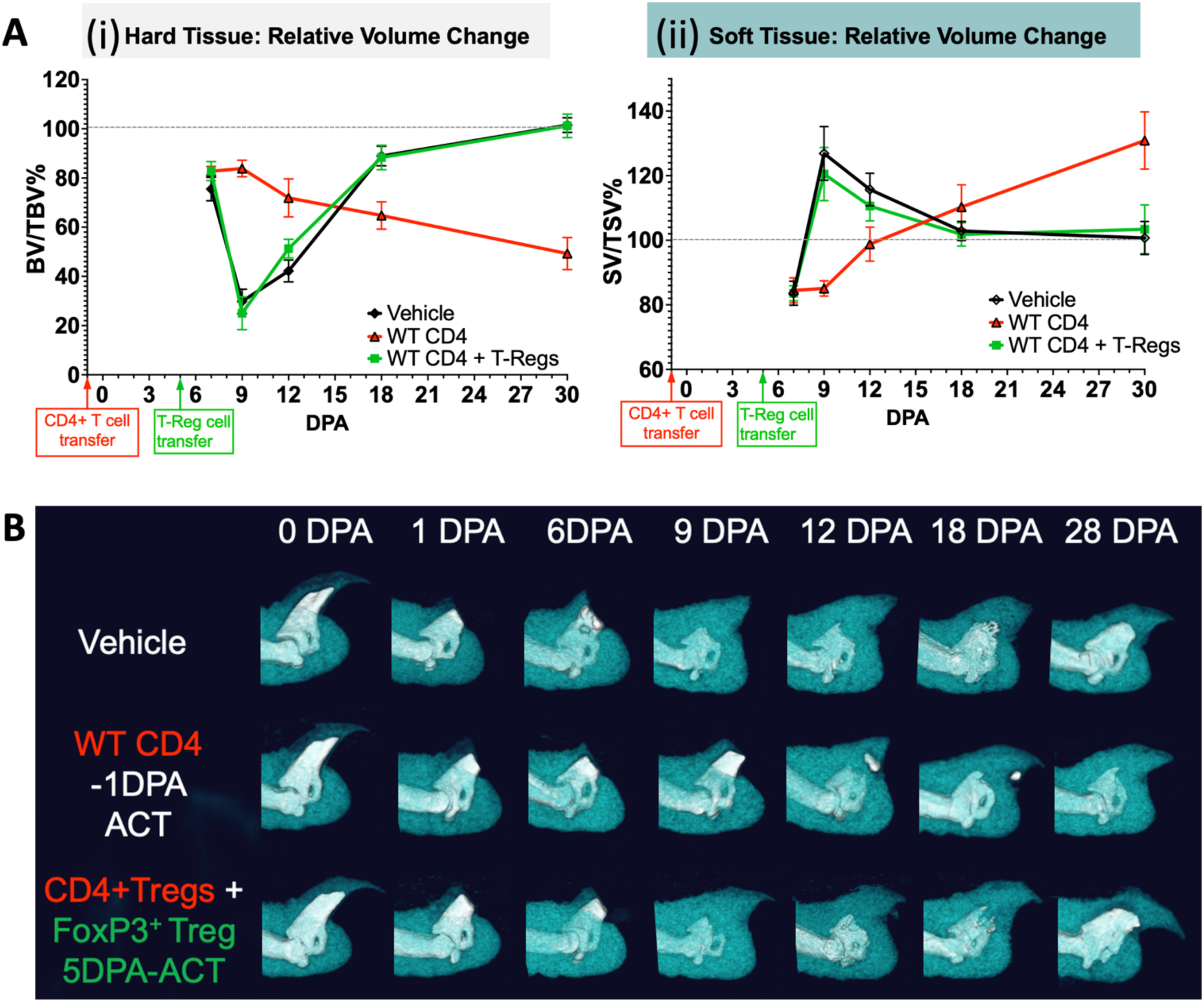
Asynchronous ACT of T-regs can rescue CD4^+^ T-cell-mediated inhibition of digit-tip regeneration in B6RG mice. Lymphoid-deficient B6RG mice receiving purified CD4^+^ T cells one day prior to P3 amputation show potent inhibition of regeneration relative to vehicle controls. ACT of 3 x 10^6^ FACS-purified FoxP3^+^ T-regs delivered via tail vein at 5 DPA (6 days post CD4^+^ T-cell delivery) completely rescued regeneration to normal control levels. **A**. Changes in hard tissue (i) and soft tissue (ii) relative to uninjured control over time. Timing of interventions indicated on X-axis. N = 5 mice with 4 amputated digits per animal = 20 measurements per condition, per timepoint. Data representative of at least 2 independent experiments. BV,= bone volume, SV = soft volume, TV = total volume. P values calculated using one-way ANOVA with Turkey’s multiple comparison test. Statistical differences relative to B6 reference strain are indicated as ^ns^P > 0.05, *P ≤ 0.05, **P ≤ 0.01, ***P ≤ 0.001, ****P ≤ 0.0001. **B**. Representative two-tone microCT images of regeneration over time comparing treatments. Bone (hard tissue) shown in white, and soft tissue shown in teal.

### CD4^+^ T-cell-mediated inhibition of regeneration is associated with apoptosis and necroptosis, both of which are attenuated in T-reg-rescued digits

CD4^+^ T cells show potent inhibition of regeneration by inhibiting the osteoclastogenesis necessary for the catabolic phase of P3 digit-tip regeneration that precedes anabolic bone growth (**Fig. 5, 6, S7, S8**) In previous work, we demonstrated that ThNK cells are cytotoxic to both OC and OB progenitor cells, both of which are required for P3 digit regeneration^16^. To determine if the inhibitory effect of CD4^+^ T cells on regeneration is mediated at least in part by cytotoxic activity against OCs, we performed ACT of CD4^+^ T cells in lymphoid-deficient B6RG and NRG mice with and without FoxP3^+^ T-reg rescue and then measured the number of apoptotic or necroptotic cells relative to vehicle control animals, via immunostaining. Results show that in both strains, the number of TUNEL^+^ apoptotic cells was significantly increased in the amputated digit at 7 DPA and was abolished via T-reg rescue (**Fig. 7A, B, Fig. S10D**). We also observed CD4^+^ T cells in close association with TUNEL^+^ apoptotic cells (**Fig. 7C**). We then assessed whether caspase-independent necroptotic death could also be taking place, by measuring phopho-RIP3, which provides a snapshot of necroptotic activation ^41^ ^42^. This analysis showed a distinctive cluster of necroptotic cells in the marrow only in the CD4^+^ T-cell condition, and an alternative pattern of necroptotic activity in the dorsal mesenchyme only in the CD4^+^ T-cell/T-reg rescue cohort **(Fig. 7D)**, suggesting that FoxP3^+^ T-reg-dependent rescue may involve anti-CD4^+^ T-cell necroptotic activity. We then measured OC numbers in the different conditions via the OC marker, cathepsin K (CathK), and observed a significant reduction in OC numbers between 7 DPA and 9 DPA in CD4^+^ T-cell-treated mice that was rescued to normal levels via FoxP3^+^ T-reg chase ACT (**Fig. 7E, F)**. This is consistent with results observed in NRG T-reg rescue of CD4^+^ T-cell inhibition using TRAP enzymatic staining to measure OC numbers (**Fig. S10C**).

**Figure 7.**
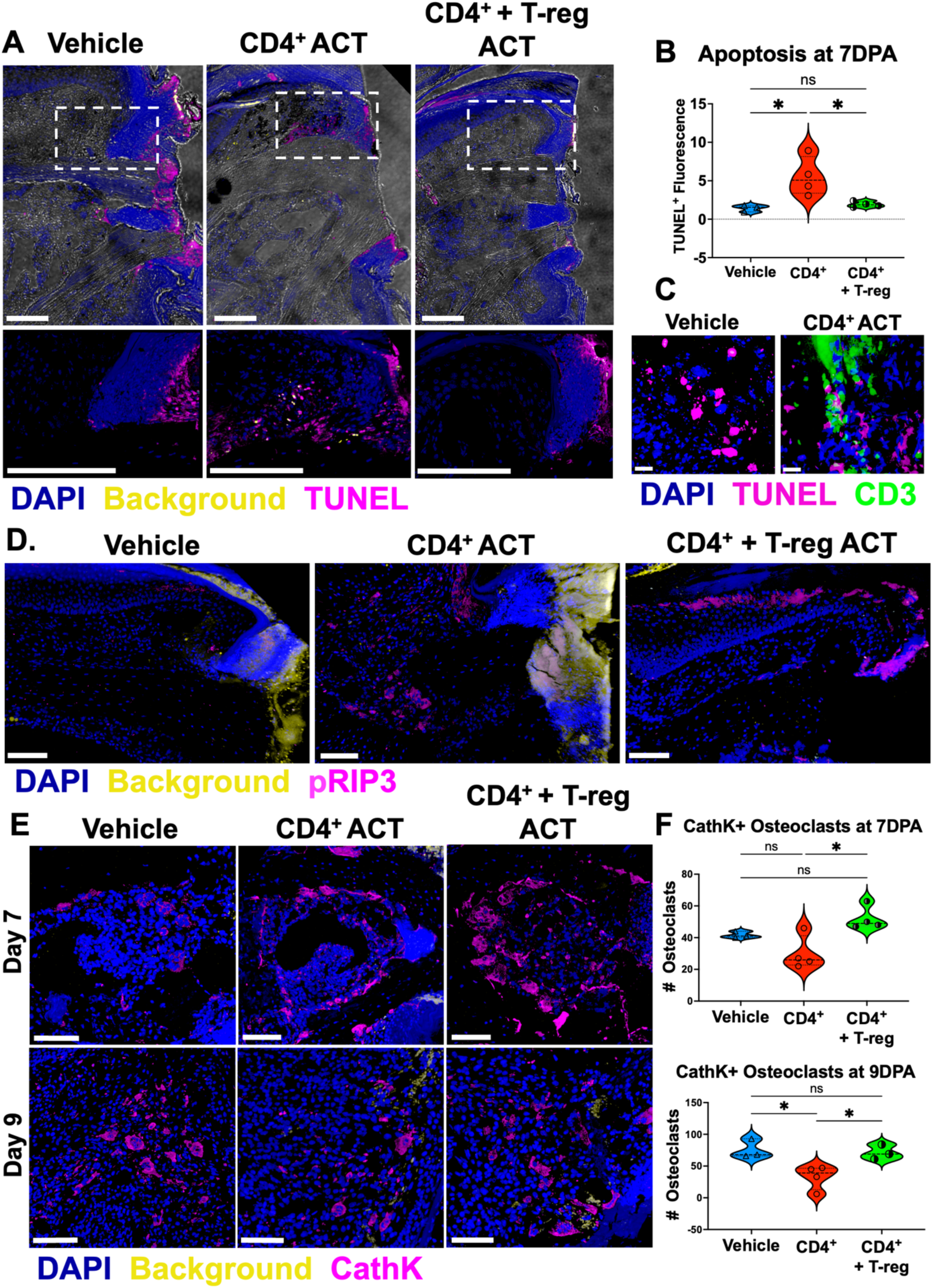
CD4^+^ T cells induce apoptosis and necrosis and inhibit osteoclast survival, all of which are rescued with asynchronous ACT of FOXP3 T-regs. **A.** TUNEL^+^ immunofluorescent labelling in B6RG and NRG mice shows an accumulation of apoptotic cells at 7 days post amputation (7 DPA) only in digits receiving CD4^+^ T-cell ACT one day prior to amputation. TUNEL^+^ cell induction is abolished upon asynchronous systemic delivery of FoxP3^+^ T-reg ACT at 5 DPA via tail-vein injection. Scale bar = 150 μm. (paraffin sections) **B.** Quantification of TUNEL^+^ cells at 7 DPA. **C.** High magnification immunofluorescence image of CD3^+^ T cells at 7 DPA closely associated with TUNEL^+^ apoptotic cells in B6RG lymphoid-deficient mice that received ACT of CD4^+^ T cells one day prior to amputation. Scale bar = 10μm. (paraffin sections) **D.** Phospho-specific RIP3 immunostaining reporting cellular necrosis is induced in CD4^+^ T-cell ACT-treated animals at 7DPA. This pattern of necrotic pRIP3 is altered via FoxP3-T-reg rescue in CD4^+^ T-cell-inhibited mice. Scale bar = 100μm. (Cryosections) **E.** Cathepsin K (CathK) immunostaining shows that CD4^+^ T-cell ACT treatment (-1 DPA) reduces osteoclast survival between 7 DPA and 9 DPA. **F.** Quantification of mature multinucleated CathK osteoclasts in each treatment group. Scale bar = 100μm. (paraffin sections) P values calculated using one-way ANOVA with Turkey’s multiple comparison test. Statistical differences relative to B6 reference strain are indicated as ^ns^P > 0.05, *P ≤ 0.05, **P ≤ 0.01, ***P ≤ 0.001, ****P ≤ 0.0001.

**Figure 8.**
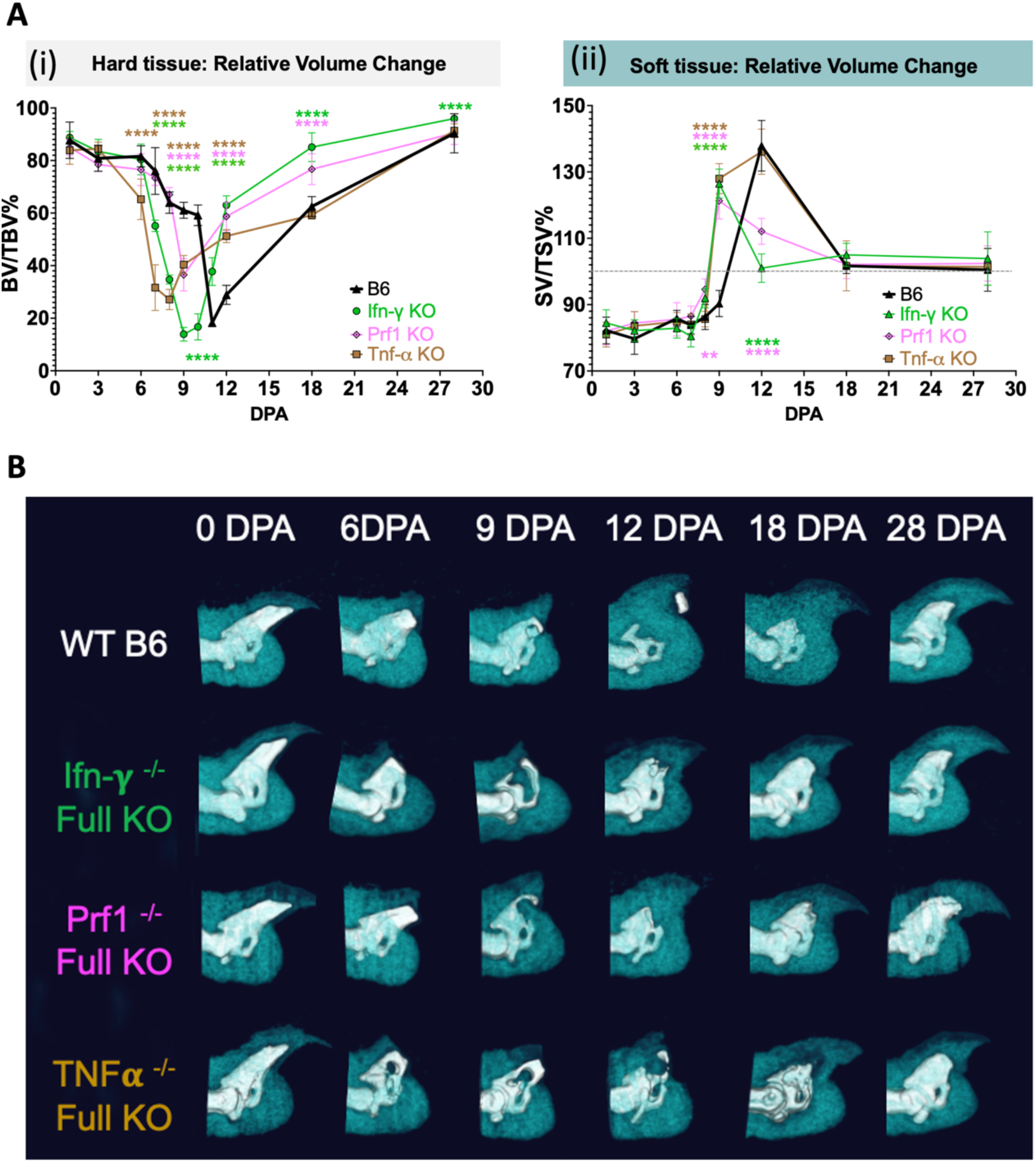
Digit-tip regeneration assays using full knockouts of perforin, IFN-γ, and TNF-α in mice on the B6 genetic background to identify potential cytotoxic mechanisms of CD4^+^ T-cell-mediated inhibition of regeneration. **A.** Changes in hard tissue (i) and soft tissue (ii) relative to uninjured control over time. N = 5 mice, with 4 amputated digits per animal = 20 measurements per condition, per timepoint. Data representative of at least 2 independent experiments. BV = bone volume, SV = soft volume, TV = total volume. P values calculated using one-way ANOVA with Turkey’s multiple comparison test. Statistical differences relative to B6 reference strain indicated as ^ns^P > 0.05, *P ≤ 0.05, **P ≤ 0.01, ***P ≤ 0.001, ****P ≤ 0.0001. Means and standard deviations are shown. **B.** Representative two-tone microCT images of regeneration over time comparing strains. Bone (hard tissue) shown in white, and soft tissue shown in teal.

### Genetic knock-out of key cytotoxicity genes (IFN-γ, Prf1, TNF-α) enhances regeneration

To gain greater insights into the mechanism of CD4^+^ T-cell-mediated inhibition of regeneration, we tested the potential participation of three major cytotoxicity genes implicated in T-cell- or NK-cell-mediated cytotoxicity. Using global genetic knock-outs of *IFN-γ, Prf-1, and TNF-α* in wildtype B6 mice, we observed enhanced P3 digit-tip regeneration with each knock-out that resulted in an increased speed of catabolism and an earlier switch to anabolism (**Fig. 8**). To test the role of *IFN-γ*, *Prf1*, and *TNF-α* specifically in CD4^+^ T cells, we performed ACT of mutant magnetically purified *IFN-γ* KO, *Prf1* KO, *TNF-α* KO or wildtype CD4^+^ T cells into B6RG lymphoid-deficient mice and compared their digit-tip regeneration to vehicle-treated mice (**Fig. 9**). Whereas Prf1-deficient CD4^+^ T cells were unable to rescue normal catabolism, loss of TNF-α in CD4^+^ T cells substantially attenuated CD4^+^ T-cell-mediated inhibition of regeneration, significantly improving the rate of catabolism and partially rescuing bone regeneration. In contrast, IFN-γ-deficient CD4^+^ T cells produced a complete rescue of regeneration to the levels seen in vehicle-injected mice, confirming a critical requirement for IFN-γ in suppression of osteoclastogenesis and catabolism necessary for P3 digit-tip regeneration.

**Figure 9.**
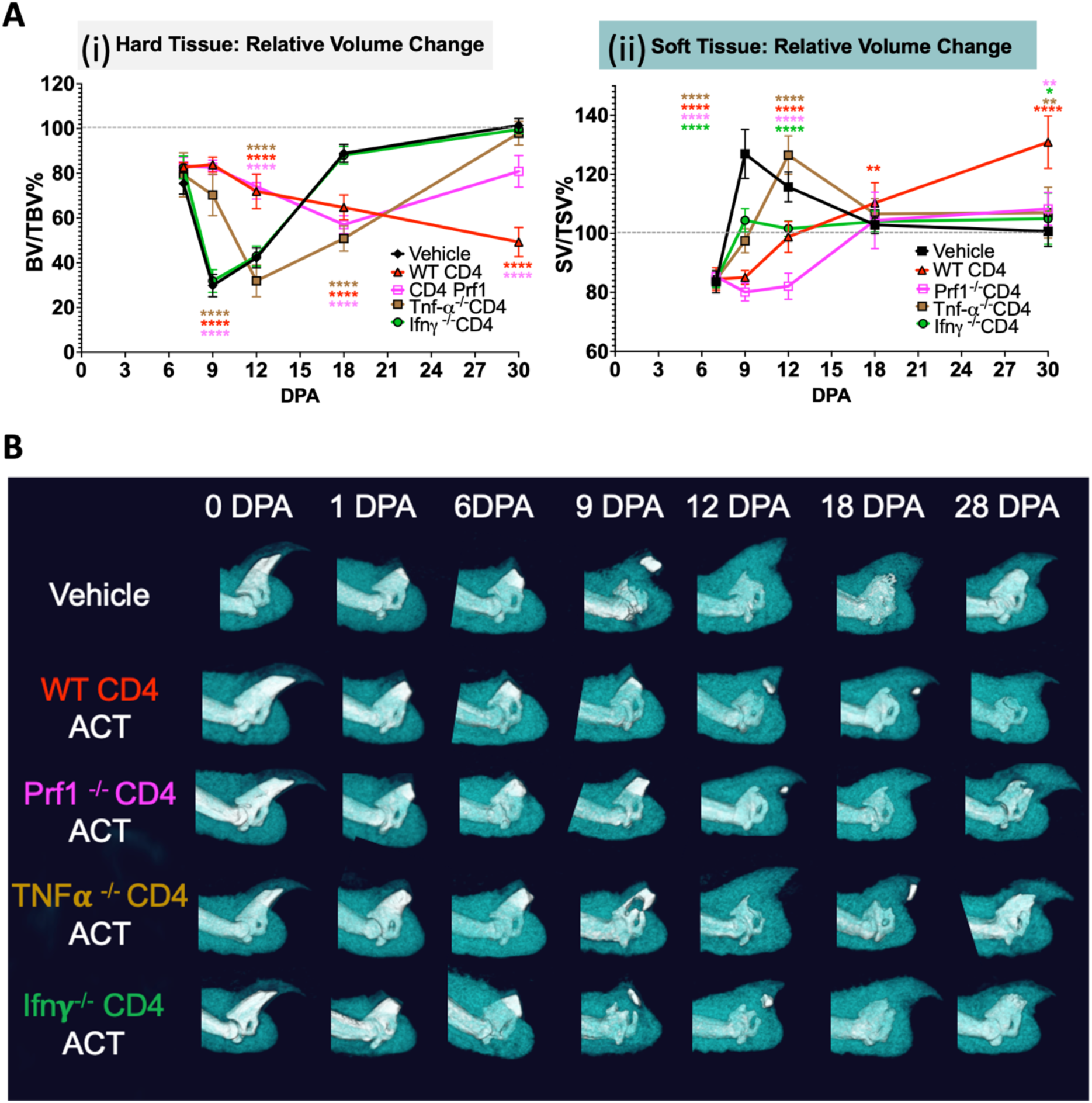
ACT of mutant CD4^+^ T cells lacking perforin, IFN-γ, or TNF-α into lymphoid-deficient B6RG mice shows that IFN-γ is the major cytotoxic effector in CD4^+^ T-cell-mediated inhibition of regeneration. **A.** Changes in hard tissue (i) and soft tissue (ii) relative to uninjured control over time. N = 5 mice, with 4 amputated digits per animal = 20 measurements per condition, per timepoint. Data representative of at least 2 independent experiments. BV = bone volume, SV = soft volume, TV = total volume. P values calculated using one-way ANOVA with Turkey’s multiple comparison test. Statistical differences relative to B6 reference strain indicated as ^ns^P > 0.05, *P ≤ 0.05, **P ≤ 0.01, ***P ≤ 0.001, ****P ≤ 0.0001. Means and standard deviations are shown. **B.** Representative two-tone microCT images of regeneration over time comparing ACT treatments with different mutant CD4+ T cells and saline vehicle treated controls. Bone (hard tissue) shown in white, and soft tissue shown in teal.

## Discussion

Regenerative capacity in the digit tip is highly conserved in mice and humans, underscoring the value of mouse digit-tip regeneration studies for clinical applications. The mouse P3 digit-tip regeneration model is one of very few well-characterized platforms for studying adult epimorphic regeneration. The present study advances our understanding of the regenerative process through several important findings: **1)** Regeneration is enhanced in the absence of all lymphoid immune cells. **2)** Naïve CD4^+^ T cells are potent inhibitors of the regenerative response, a capacity that we previously demonstrated for thymic-derived NK cells. **3)** FoxP3^+^ T-regs can enhance the OB-dependent anabolic phase of regeneration in the absence of other lymphoid cells. **4)** FoxP3^+^ T-regs can block the inhibitory role of CD4^+^ T cells, restoring normal osteoclastogenesis, and are necessary for normal regeneration in immunocompetent animals. **5)** The mechanism of CD4^+^ T-cell-mediated inhibition of regeneration involves IFN-γ-mediated induction of apoptosis and necroptosis. **6)** RANKL-mediated osteoclastogenesis is a rate-limiting step for digit-tip regeneration, and CD4^+^ T-cell-mediated inhibition of regeneration can be overcome with recombinant RANKL supplementation. Collectively, these results significantly increase the potential the mouse digit tip model for promoting therapeutic regeneration in humans.

### Fracture healing and digit tip regeneration compared

This work reveals that the osteoimmunology of the mammalian digit tip is a key determinant of regeneration success and provides the first comprehensive assessment of the role major T cell subset roles in digit tip regeneration. These results must be contextualized with the current osteoimmunology knowledge in fracture repair despite epimorphic digit-tip regeneration involving different processes. In fracture repair, inflammation precedes and regulates the immune-cell recruitment and OC formation required to dissolve and remodel damaged bone^43^. New soft callus composed of cartilage and fibrous tissue is laid down via chondrocytes to bridge the fracture gap, where OBs are activated to carry out endochondral ossification, i.e., replacement of the soft callus with hard callus composed of bone that undergoes subsequent mineralization and ossification. Subsequently, a lengthy remodeling process involving OCs and OBs convert the newly formed and relatively loosely organized woven bone into highly organized mature lamellar bone. In contrast, P3-level digit-tip amputations (distal 1/3 removed) in mammals regenerate through direct intramembranous ossification without cartilage intermediates but by forming a mound of undifferentiated cells known as a blastema. In this alternative mode of repair, only around 15% of the P3 bone volume is lost initially, but the inflammatory response and immune-cell recruitment induce robust OC-driven catabolism of the P3 bone well below the amputation plane, sometimes leaving behind as little as 20% of the original bone volume, making space for the digit-tip blastema. OBs are then induced that directly lay down new woven bone that is ossified, and then, after the digit is fully regenerated, the new bone undergoes remodeling to generate stronger lamellar bone. One major advantage of studying bone repair in the digit tip model is the greatly simplified and enhanced sensitivity of measuring catabolic and anabolic phase kinetics. Although fracture healing forms fibrocartilaginous tissue that is not present in digit tip blastema, both models restore bone through some shared mechanisms that rely on finely tuned interactions between skeletal and immune cells.

### The impact of T and B cells on bone quality in fracture healing and regeneration

Many studies have tested T cell function in fracture healing models but have not reached consensus. One fracture healing study using RAG1-KO mice, lacking T and B cells, demonstrated accelerated regeneration of hard and soft tissues^44^ similar to what we observed in the mouse digit tip. Rag1-KO mice exhibit earlier soft callus cartilage mineralization and remodeling after fracture, suggesting improved regeneration in the absence of T cells. Other fracture studies have suggested that rapid endochondral ossification and mineralization caused by the lack of T and B cells may have a deleterious impact on healing by generating higher stiffness and decreased bone quality^45^. This raises the question of whether the lack of T cells in our digit-tip regeneration model affects the quality of new bone. However, digit tip regeneration includes neither cartilage formation nor endochondral ossification, the latter of which appears to be the major step affecting bone quality during fracture repair RAG-1-KO mice. Furthermore, although we have previously defined the role of NK cell in digit tip regeneration^16^, the role of NK cells in fracture healing is poorly defined and RAG-KO mice retain hyperresponsive NK cels^46^ ^47^ that are affected by the absence of T and B cells and the RAG deletion effects on NK-cell heterogeneity and fitness^46^. In the digit tip we found that regeneration was even faster in mice with both RAG-KO and IL2-Rγ KO, the latter of which eliminates development of NK cells. Our fastest regenerators were the two nude strains Nu/J and B6:FoxN1, both of which lack T cells, B cells, and thymic NK cells, but carry splenic NK cells. These results are consistent with our previous findings of a pro-regenerative splenic NK-cell population and an inhibitory thymic NK-cell population with potent cytotoxic effects on OC and OB progenitor cells^16^. More broadly, previous studies are inconclusive regarding whether T and B cells exert a net positive or negative effect on bone health or repair. For example, activated T cells have been shown to cause bone loss during rheumatoid arthritis^48^ and postmenopausal osteoporosis^49^, and CD8^+^ memory/effector T cells have been linked to delayed bone healing following fracture in humans^19^. Taken together, it is clear that bone repair is strongly impacted by immune-cell repertoire.

### The role of T cells in other contexts of regeneration and development

The role of T cells in repair and regeneration is complex. Reflecting this complexity, studies have yielded results that differ depending on the context. Results of a skin-repair model suggested that CD4^+^ T cells are the predominant lymphocyte in the dermis of hypertrophic scars, the layer where some of the lymphokines act on keratinocytes, fibroblasts, and other cell types to induce fibrosis ^50^. However, historical wound-healing studies have shown conflicting results regarding whether T cells promote or inhibit healing. Antibody depletion of CD4^+^ and CD8^+^ T cells in rats showed delayed skin healing following thymectomy^51^ ^52^. In contrast, work using mice lacking either CD4^+^ or CD8^+^ T cells suggested that the absence of either cell type had no effect on wound closure, wound-breaking strength, collagen content, or angiogenesis^53^. In a model of skin injury, nude mice, which lack T cells, B cells, and thymic-derived NK cells, showed altered scar mechanotransduction compared to wildtype mice, characterized by altered recruitment of macrophages and fibroblast precursors, and altered epidermal and dermal proliferation, in response to mechanical loading^54^. These results support our findings using T-cell ACT in nude mice, showing that T cells impact myeloid-cell phenotypic diversity. A study investigating the role of T cells in cardiac injury and repair showed that RAG-1-KO mice exhibit reduced infarct size after cardiac injury, and that this reduced size can be increased to the infarct size in wildtype B6 mice after adoptive transfer of wildtype CD4^+^ T cells but not IFNγ-KO T cells^55^, suggesting that the IFNγ-dependent cytotoxic effects of CD4^+^ T cells on progenitor cells that we observed in our digit-tip model may be a generalized phenomenon and not bone-specific. Consistent with this notion, a study using mouse mammary-gland (MG) organoids showed that IFN-γ secreted by CD4^+^ T cells disrupts epithelial branching morphogenesis in MG organogenesis^56^. In other work, depletion of CD4^+^ T cells could not promote post-injury regeneration in the adult mouse heart but *<u>could</u>* enhance post-injury repair in the non-regenerative neonatal mouse heart (postnatal day 8), with the enhanced repair linked with promotion of cardiomyocyte survival and attenuation of fibrosis^57^. These results suggest that early postnatal T-cell repertoires and those of adult tissues may differ significantly.

T regulatory cells have been shown to be essential for zebrafish cardiac regeneration^58^. Similarly, T-regs are necessary for mouse neonatal (postnatal day 3) cardiac regeneration, by supplying paracrine factors to promote cardiac proliferation, and by shaping macrophage polarization to attenuate fibrosis^59^. In addition, a study using spiny mice (*Acomys* spp.) showed a role for T-regs in regeneration. Spiny mice, which are more closely related to gerbils than to laboratory mice^60^, have emerged as a powerful mammalian model of scar-free repair and epimorphic regeneration. *Acomys cahirinus* was shown to exhibit a robust T-cell response rich in genetic signatures for T-regs and low for CD4^+^ T-cell signatures relative to *Mus musculus* (the outbred Swiss Webster strain)^61^. Together, these studies are consistent with a positive role of T-regs and a negative role of CD4^+^ T cells in epimorphic regeneration.

Our experiments in the skin injury model, i.e., nude mice, which lack T cells, B cells, and thymic-derived NK cells, suggests that the phenotypes of myeloid cells recruited to skin wounds differ depending on the presence or absence of adoptively transferred T cells, a finding that is supported by work from Wong et al.^54^. However, a limitation of our experiment is that immunophenotyping in the skin model may not completely reflect immunology of the digit tip. Further, the flow-cytometry approach we used for immunophenotyping in the skin model is technically challenging because of the very high cell input required. Future studies using single-cell RNA sequencing approaches allowing low cell inputs would be useful for testing whether T-cell restriction of myeloid heterogeneity extends completely to the digit tip.

Our experiments show that CD4^+^ T cells activate progenitor-cell apoptosis and thereby inhibit the rate-limiting step of new bone regeneration, i.e., differentiation of monocyte/macrophage progenitor cells into OCs, consequently blocking new bone growth. We also showed that this blockage of new bone growth can be completely rescued by a chase with adoptively transferred FoxP3^+^ T-regs, which directly suppress the CD4^+^ T-cell activity. T-regs have also been shown to inhibit IFN-γ-/TNF-α-dependent T-cell-mediated destruction of transplanted bone marrow mesenchymal stem cells (BMMSCs)^62^, supporting our results. Concordantly, studies have shown that CD4^+^ T cells and CD8^+^ T cells negatively regulate blood-vessel regeneration, and FoxP3^+^ T-regs positively regulate blood-vessel regeneration^63, 64^. Because the digit-tip blastema is avascular at the above rate-limiting step in bone regeneration, CD4^+^ T-cell-mediated inhibition of angiogenesis is unlikely to play a major role in this early phase of regeneration. However, given that monocytes/macrophages direct new vessel development^65^ and that, as we show above, CD4^+^ T cells can shape myeloid-cell phenotypes, CD4^+^ T cells may play an as-yet-undefined role in the later anabolic phases.

### The impact of IFN-γ on regeneration

Although in this study IFN-γ expression by CD4^+^ T cells was required for inhibition of osteoclastogenesis via progenitor apoptosis, the effects of IFN-γ on OC formation are controversial in other models. It is likely that the effects of IFN-γ on OC formation are complex and are influenced by particular conditions in the bone microenvironment. For example, IFN-γ is a strong suppressor of osteoclastogenesis *in vitro* through inhibition of RANKL signaling^66^, but enhances OC generation in cultures of peripheral blood from osteoporotic patients^67^. *In vivo* studies in three mouse models of bone loss (ovariectomy, LPS injection, and inflammation induced by silencing TGF-β signaling in T cells) contradicted our findings, showing that the net effect of IFN-γ in these conditions was bone loss due to increased OC formation through T-cell expression of RANKL and TNF^68^. Other studies are consistent with our observations, showing that IFN-γ directly inhibits TNF-α-induced osteoclastogenesis *in vitro* and *in vivo* and induces apoptosis mediated by Fas/Fas ligand interactions^69^. Similarly, *in vivo* work in mice showed that IFN-γ released by CD4^+^ T cells induced apoptosis of BMMSCs^70^.

### Role of osteoclasts in bone-tissue integration

In the salamander limb regeneration model, OC-driven bone resorption is required for successful integration of the new regenerated bone with the existing stump ^71^. In a mouse study, OC inactivation did not impact OB bone formation but negatively impacted fracture healing by delaying the resorption of cartilaginous callus and consequently impeding the remodeling of immature bone to mature lamellar bone and impairing bone biomechanical strength^43^. Concordantly, mice lacking osteoprotegerin, a protein that decreases OC activity, exhibited accelerated resorption of the cartilaginous callus and significantly enhanced union of fractured bone^72^. Another study showed that mice lacking DAP-12, a protein involved in induction of OC-specific genes^73^, showed a decrease in cartilaginous resorption and in formation of woven bone, as well as bone structural defects^73^. A pharmacological study in rats showed that inhibition of OC formation and/or bone resorption delays fracture healing by disrupting cartilage dissolution and remodeling, and increases fracture callus size^74^.

In the mouse P3 digit-tip regeneration model, a single injection of free clodronate (F-Clo) immediately following amputation eliminates OCs in the P3 digit, causing severe delays in bone catabolism and bone anabolism ^75^ despite the very short half-life of F-Clo, i.e., 1-2 hours^76^. Interestingly, induction of early epidermal closure in the mouse P3 digit-tip model using a wounding glue can induce direct lamellar bone extension, bypassing the normal OC-mediated catabolism-to-OB-mediated anabolism program that results in formation of woven bone^77^. Based on the requirement for bone resorption in both salamander limbs stump ^71^and fracture healing^43^ ^73^ one could predict that bypass of this program, including normal OC function, would result in inefficient integration of new bone and, consequently, a final bone that is potentially more prone to fracture. However, the biomechanical properties of bone formed via this approach have not yet been evaluated.

This study has shed new light on the role of different major T cell types and revealed a mechanism of regulating progenitor cell survival through CD4+ T cell and FoxP3+ T cell antagonism. Although this work identified that the main mechanism of CD4^+^ T-cell-mediated inhibition of regeneration is IFN-γ-mediated cytotoxicity, IFN-γ expression levels may differ among different T-cell subsets differentiating from naïve CD4 cells (e.g., T-reg, Th1, Th17, Th2, T_FH_, Th9, and Th22)^17^. Although it is difficult to assess *in vivo* differentiation of splenic naïve CD4^+^ T cells into the various T-cell subsets, one can predict, based on the fact that IFN-γ is among the cytokines associated with classic Th1 or Th17 cells^17^, that naïve CD4^+^ T cells may differentiate into these two CD4^+^ T-cell subsets in the digit tip. Other CD4^+^ T-cell subsets (i.e., Th2, Th9, and Th22) are not usually associated with significant IFN-γ expression^17^, but more work is required to thoroughly understand differentiation of naïve T cells recruited to the amputated digit tip.

## Conclusions

Innate and lymphoid immune cells are components of a dynamic network that prepares the injury site for progenitor-cell mobilization and productive regeneration. Successful formation of the blastema likely demands a very precise inflammatory milieu. An overzealous immune response, particularly from proinflammatory T-cell subsets, disrupts the dedifferentiation of local cells or leads to scarring, thereby negating the regenerative process. Conversely, a threshold of T-regs and other supportive immune signals might be necessary to prime the tissue for re-growth. Regenerative success relies on a delicate balance between inflammatory pathways and the cellular crosstalk that control survival and proliferation of the progenitor cells required for each phase of regeneration. The present work has identified a cytotoxic CD4^+^ T-cell subset that can kill endogenous digit-tip progenitor cells via a mechanism distinct from that used by thymic NK cells, which exert their cytotoxic effects via a Perforin/Granzyme mechanism^16^. We also showed that these cytotoxic effects of CD4^+^ T cells can be controlled via supply of T-regs or by CD4^+^ T-cell-specific IFN-γ deletion. We found that supply of excess T-regs has no effect on the catabolic phase of regeneration but enhances OB-driven bone growth. However, we also showed that selective T-reg ablation destroys the T-cell balance, allowing inhibition of osteoclastogenesis to dominate, in turn delaying regeneration. Taken together, our work clearly demonstrates that lymphoid immunity has a net negative influence on digit-tip regeneration but can be enhanced with T-reg supplementation. Understanding the key regulators and cellular networks controlling each stage of regeneration will ultimately lead to new insights and approaches to expand the regenerative capacity of human patients.

## Materials and Methods

### Mice

Mice were acquired from The Jackson Laboratory (JAX; Bar Harbor, Maine) and housed in a pathogen-free facility on a 12:12 hour light:dark cycle on normal chow. Male mice 10-16 weeks old were used for all studies and were aged-matched for all strains within each experiment. All animal experiments were carried out according to procedures approved JAX’s Animal Care and Use Committee (IACUC) and comply with all relevant ethical regulations regarding animal research. Strains used in this study are listed below.

(B6) C57BL/6J, Strain #000664,
(mT/mG) Membrane-TdTomato mice, (B6.129(Cg)-Gt(ROSA)26Sortm4(^ACTB-tdTomato,-EGF^ ^P)Luo/J^) Strain #7676
(Athymic Nude) NU/J - Strain #002019
(BALBc) BALB/cJ- Strain #000651 (control strain for NU/J)
(B6-athymic nude), B6.Cg-*Foxn1^nu^*/J- Strain #000819
(B6 Rag1 KO), B6.129S7-*Rag1^tm1Mom^*/J- Strain #002216
(B6 Rag2 KO), B6.129S7-*Rag1^tm.1Cgn^*/J- Strain #008449
(NRG), NOD.Cg-*Rag1^tm1Mom^ Il2rg^tm1Wjl^*/SzJ - Strain #007799
(FoxP3:DTR-GFP), B6.129(Cg)-*Foxp3^tm^*^3^*^(DTR/GFP)Ayr^*/J- Strain #016958
(FoxP3:YFP) B6.129(Cg)-*Foxp3^tm^*^4^*^(^YFP^/icre)Ayr^*/J- Strain #016959
(Ncr1:GFP/ NKp46^GFP^ ^knock^ ^in/knock^ ^out^) B6;129-Ncr1^tm1Oman^/J, Strain #022739
(B6 IL2Rγ null) B6.129S4-*Il2rg^tm1Wjl^*/J- Strain #003174
(Prf1KO), C57BL/6-Prf1^tm1Sdz/^J Strain #002407
(IFN KO)B6.129S7-Ifng^tm1Ts/^J Strain #002287
(TNFα KO) B6.129S-Tnf^tm1Gkl/^J Strain # 005540
“BRG” mice (B6_RAG-KO_IL2Rγ-KO) were generated by intercrossing B6.129S7-*Rag1^tm.1Cgn^*/J (B6 Rag2 KO) with (B6 IL2Rγ null) to combine the RAG-2-KO allele and the IL2Rγ-KO allele on the C57BL6/J genetic background. Both alleles were bred to homozygosity confirmed by JAX transgenic genotyping services. KO = Knock-out allele.

### Digit-tip surgery

Digit-tip amputation experiments were performed as described previously using aged-matched 10-16-week-old mice that were anaesthetized and underwent amputation of the distal one third of the terminal phalanges^78^. Mice were anesthetized with isoflurane. We amputated two digits per rear paw (digits 2 and 4) of the hindlimb, keeping digit 3 as an uninjured control. After amputation, the digits were sprayed with Cavilon No-Sting Barrier Film (3M) to minimize potential infections.

### Microcomputed tomography (microCT) imaging, image processing, and quantification

To measure changes in regeneration phenotypes, P3 digits were scanned live at various timepoints using the Quantum GX MicroCT (Perkin Elmer) under isoflurane anesthesia. Digits were scanned at high resolution (voxel size 10μm and energy of 55 kVp; 1,000 projections per 180° captured at 380 msec using continuous rotation). MicroCT files were exported as both VOX files and DICOMS.

DICOMS were loaded into Bruker CTAn software version 1.18 with the lower limit set to 3100 to eliminate background noise. Region of interests (ROIs) were created for each single digit with the Z axis set at the boundary of P3/P2 joint. 3D volumes of hard and soft tissue were calculated with the morphometric analysis tool, setting the hard-tissue threshold at 65-165 and soft-tissue threshold at 165-255. 3D volumes were tabulated, and relative values were calculated by dividing the experimental-digit measurement by the control uninjured digit measurement (with the two digits from the same paw) expressed as a percentage (i.e., BV/TBV or SV/TSV). Data was visualized in GraphPad Prism (10.2.0) as a line graph.

To generate the “two-tone” (white bone/teal soft tissue) 3D rendering of digits at different timepoints after amputation in each cohort, representative digits falling in the mean of each group were selected for visualization. VOX files were imported into Perkin Elmer Database (version 3.5.4.110). Hard bone was assigned the color white with a threshold of >268, and soft tissue was assigned the color teal with a threshold of 99 to 268. Smooth filter 3D (-3) was turned on and the images exported as 2D BMPs for figure generation.

### Magnetic purification of leukocytes for adoptive transfer

For CD3^+^ T-cell, CD4^+^ T-cell vs. CD8^+^ T-cell adoptive transfer experiments, “untouched” cells were isolated via negative magnetic selection to prevent activation. MojoSort Mouse CD3^+^ T-cell isolation Kit (Biolegend, Cat#480031), MojoSort Mouse CD4^+^ T-cell isolation Enrichment Kit (Biolegend, Cat# 480033) or MojoSort Mouse CD8^+^ T-cell Enrichment Kit (Biolegend, Cat# 480008) without RBC lysis, was used according to manufacturer’s instructions. Adoptive transfer of leukocytes into T-cell-deficient hosts was performed via tail vein injection. Lymphoid-deficient animals were reconstituted with cell numbers aligning with their relative numbers in wildtype B6 mice (i.e., ACT was performed using 2.5 x10^7^ Pan CD3^+^ T cells, or 1.8×10^7^ CD4^+^ T cells, or 0.7×10^7^ CD8^+^ T cells, or 1 × 10^7^ NK cells, or 3 ×10^6^ FoxP3^+^ T-regs, with an average of 1× 10^8^ total leukocytes from donor spleens.

To validate the purification of magnetically enriched leukocytes, we performed adoptive transfer flow cytometry prior to adoptive transfer, for each purification. Typical purifications were over 85%-95% pure **(Fig. S6).** Single-cell suspensions were treated with unlabeled Fc receptor blocking antibody cocktail (CD16/CD32) in 50 µl of FACS buffer (HBSS + 5mM EDTA + 2% FCS). Cells were then incubated in a “staining antibody cocktail” with antibodies against surface markers including CD45 (pan-leukocyte), CD3 (T cells), CD19 (B cells), CD11b (myeloid cells), Nk1.1 (NK cells), and CD4 (T cell/DC) /CD8 (T cell/DC) subsets for 60min at 4°C in 100 µl. Cells were then washed in 1 ml of FACS buffer and centrifuged at 400g for 7min, 2 times. Cells were stained with 4’,6-diamidino-2-phenylindole (DAPI) to gate live events and analyzed on a five-laser 30-parameter FACSymphony A5 cytometer (BD Biosciences, San Jose, USA) using DAPI exclusion for cell viability. Antibodies used were from Beckton Dickinson (BD): 564279-Anti-Mouse CD45-BUV395 (30-F11), BD-565076-Anti-Mouse CD19-BUV661 (1D3), BD-553061-Anti-Mouse CD3e-FITC (145-2C11), BD-563068 Anti-Mouse CD8a-BV510 (53-6.7), BD-552051-Anti-Mouse CD4-APC-Cy7(RM4-5), BD-564443-Anti-Mouse CD11b-BUV737 (M1/70), and BD-553165-Anti-Mouse NK1.1-PE (PK136). Data was analyzed with FlowJo version 10 (Tree Star, Ashland, OR).

To isolate untouched T-regs for adoptive transfer, FoxP3-YFP spleens were surgically removed, pooled, placed in ice cold HBSS+2% Fetal Calf Serum (FCS) before manual dissociation through 100 µm sterile mesh filters. Cells were washed in 20 ml of HBSS+FCS and pelleted at 200 x g for 20 minutes in a 50 ml conical tubes. Spleen cells were immersed in 10 ml of AKC lysis buffer (150 mM NH_4_Cl, 10 mM KHC0_3_, 0.1 mM Na_2_EDTA, pH 7.4), mixing gently. Cells were incubated at room temperature for 10 minutes before being centrifuged at 400 x g for 5 minutes. Pelleted cells were resuspended in HBSS +2% FCS, and YFP^+^ T-regs were recovered to 100% purity using a FACSymphony instrument, resuspended in sterile PBS, and injected into recipient mice via the tail vein.

### *In vivo* antibody blockade of RANKL

To neutralize RANKL and block RANK-RANKL signaling *in vivo*, we used 4 doses of *in vivo*-grade blocking antibody against RANKL or isotype control via intraperitoneal (IP) delivery of 100 µg at day 6 DPA, 8 DPA, 10 DPA and 12 DPA. To block RANKL, InVivoMAb anti-mouse RANKL (CD254) (Clone: IK22/5) (Bio X Cell, #BE0191) Rat IgG2aκ was used and compared against the isotope control antibody InVivoMAb rat IgG2a isotype control (Bio X Cell BE0089).

### Recombinant RANKL delivery to mice *in vivo*

Delivery of soluble RANKL into mice *in vivo* was performed by injection of 5 µg of recombinant mouse RANKL (enQuire BioReagents) into the fat pad on each hind paw at 6 DPA followed by a maintenance dose of 5 µg IP at 8 DPA and 10 DPA.

### *In Vivo* depletion of T-regs

FoxP3-DTR mice were depleted of FoxP3 T-regs by i.v. injection of 40 ng/g diphtheria toxin (DTx; Sigma) dissolved in PBS, (i.e., 1 µg in 100 µl for a 25 g mouse), starting 1 day prior to amputation and then dosed every 3 days for the duration of the experiment to maintain depletion. Vehicle controls performed in parallel.

### Immunostaining of mouse digit tips

For immunostaining, mice were perfused in PBS and digit tips were then harvested and fixed in zinc-buffered formalin, (Z-fix, Anatech Ltd) for 24 hours before decalcification in either Surgipath decalcifier I (Leica) for 48 hours at 4°C, or for 14 days in 10% EDTA. For paraffin embedding, digits were washed in PBS, transferred to 70% ethanol and then embedded in paraffin via an automated embedding processor. Five-μm paraffin sections were cut on a microtome and placed on Superfrost Plus slides (Fisher) and then air-dried overnight. Sections were deparaffinized through a xylene and descending ethanol series. Antigen retrieval was performed using Aptum R-Universal retriever buffer (#AP0530-500) and the Aptum 2100 Antigen Retriever (R2100-US) as per manufacturer instructions. Sections were permeabilized with 0.1% Triton X (Sigma T8787) for 10 minutes, washed and blocked in 5% goat serum for 30 minutes at room temperature before staining with primary antibodies (antibody details provided below). Washes were performed with PBS containing 0.05% Tween (Sigma: 655204) between primary and secondary antibody but washed in PBS several times before treatment with TrueBlack Plus (Biotium #23014) to quench autofluorescence. Sections were counterstained with DAPI for 10 minutes at 5 µg/ml before mounting in Vectashield Vibrance (Vector Laboratories H1700) mounting medium. CD68, PCNA and PHH3 images were taken on a widefield Zeiss Colibri 5-channel LED microscope using Zen Blue with single bandpass configuration. Images taken at using EC-Plan-NeoFluar 10x/0.30 M27 using an Axiocam 705 camera (2464×2056pixels). Phospho-histone H3 (Invitrogen, # 44-1190G), PCNA (DAKO, M0879) were stained at 1:100 overnight at 4°C. Detection was performed using Goat anti-Rabbit 647 (Invitrogen, A21244) and/or Goat anti-Mouse-568 (Invitrogen, A11031). Differences between treatments were determined using a one-way ANOVA with Tukey post hoc (GraphPad Prism 10.2.0).

To measure TUNEL-positive cells, 5-µm paraffin sections were deparaffinized and stained following manufacturer’s instructions using the Click-iT Plus TUNEL Assay for In Situ Apoptosis Detection, Alexa Fluor 594 or Alexa Fluor 647 (Invitrogen, C10625). CD3 staining was performed in parallel sections with anti-mouse CD3 conjugated to PE/Cy5 (1:50; Biolegend 100309) overnight at 4°C. TUNEL and CD3 imaging was performed using a Leica Stellaris S8-WLL Stellaris at 20x (NA=0.75_Glyc) utilizing the TauSense fluorescence lifetime-based TauSeperation setting^79^ to separate the unwanted background and erythrocytes with two digital gates at 0.4 ns and 1.65 ns. All images also had additional background removal using an empty 488-nm laser channel. Imaging parameters were identical for all samples. Imaging processing was done using Imaris 10.0 (Oxford Instruments) and presented as 3D projections with the DIC channel transparent for landmark identification in some cases to help with landmark identification. To determine mean fluorescence intensity of TUNEL staining in the upper mesoderm, an area approximately 300 µm in length between the nail and bone, starting behind the behind the scab and extending proximally (vehicle n=3; CD4^+^/CD4^+^ T-reg n=4). The full thickness of the section within this area was analyzed for mean fluorescence intensity using Imaris 10.0 (Oxford Instruments). Differences between treatments were determined using a one-way ANOVA with Tukey post hoc (GraphPad Prism 10.2.0).

To determine the effect of T-cell adoptive transfer on osteoclast number at 7 DPA and 9 DPA, 5-µm paraffin sections were deparaffinized and antibody-stained overnight at 4°C with anti-cathsepsin K (1:100; Abcam ab188604) and Alexa Fluor Plus 647 Goat anti-Rabbit (1:500; Invitrogen # A32733). To quantify necroptosis using RIP3 activation, a phospho-RIP3 antibody (Thr231/Ser232)(Cell Signaling; E7S1R, Cat# 91702) was applied to 7 DPA cryosections that had been fixed in 4% PFA, decalcified in 10% EDTA for 14 days, and cryopreserved in 15% sucrose/15% cold water fish gelatin for 24 hours before embedding in O.C.T. Five-µm sections were stained at 1:100 overnight at 4°C. pRIP3 was detected with Alexa Fluor Plus 647 Goat anti-Rabbit (1:500; Invitrogen # A32733). CathK and pRIP3 slides were then mounted and imaged as described above using the Leica Stellaris with TauSeperation at 20x (NA=0.75 HC PL APO_Glyc). Images were 3D-projected using Imaris 10.0, and then all individual osteoclasts in the section were counted (n=3-4). Differences between treatments at 7 DPA and at 9 DPA were determined using a one-way ANOVA with Tukey post hoc (GraphPad Prism 10.2.0).

To show that red fluorescent labelled T cells were recruited to the amputated digit post adoptive transfer, 8-µm cryosections were rehydrated in PBS, nuclear-stained with Reddot2 (1:200, Biotium #40061), and mounted using VectaShield Vibrance Mounting Media. Endogenous tdTomato expression was visualized using a Leica DIVE-4Tune/FALCON 2P with MaiTai HP (tdTomato and bone, 1030 nm) and Insight X3 (Reddot2, 1250 nm) lasers sequentially with FLIM. Bone autofluorescence was captured using a secondary detector.

### TRAP^+^ osteoclast and Masson’s trichrome staining

Paraffin sections were prepared as previously described. To measure TRAP^+^ osteoclasts, we used a protocol similar to that of Shem et al. ^80^. Briefly, after deparaffinization, sections were incubated in acetate-tartaric acid buffer (0.2 M sodium acetate (Merck, Catalog Nr. 6268), 50 mM tartaric acid, (Sigma-Aldrich, Catalog Nr. T10-9) pH 5.0) for 20 minutes at 37^°^C before incubation in 1.1 mg/ml FAST Red TR (Sigma, F8764-16) substrate and 0.1mg/ml-0.5 mg/ml napthol ASMX phosphate (Sigma N4875) substrates. Sections were incubated for 2-4 hours until sufficient red color was observed in osteoclasts. Sections were then washed and counterstained with methyl green counterstain (Vector Laboratories, H-3402-500) before washing, air drying, and mounting in VectaMount permanent mounting medium (Vector Laboratories, Cat# H-50000). Masson’s trichrome staining was performed using the Masson’s Trichrome Stain Kit SKU# KTMTRLT EA, StatLab. TX)

### Statistical information

Statistical analysis was performed using GraphPad PRISM 10.2 (GraphPad Software, La Jolla, CA). P values were obtained using either one-way ANOVA with Tukey’s multiple comparison test, or by a two-tailed unpaired t test as indicated in legends. Error bars are represented as mean ± SD. Statistical differences between groups were calculated with the following significance levels: ^ns^P > 0.05, *P ≤ 0.05, **P ≤ 0.01, ***P ≤ 0.001, ****P ≤ 0.0001. Coding of samples and blinding was done wherever possible. Rate-of-change graphs were generated by calculating the first derivative for individual curves. Means and 95% confidence intervals were assessed by one-way ANOVA with Tukey’s multiple comparison tests.

### Back biopsy model and immunophenotyping

Purified CD3^+^ T cells were isolated via magnetic negative selection from BALB/C mice for adoptive transfer into nude mice recipients. Donors and recipients were age matched. All cells were transplanted by tail vein injection. We transplanted 1× 10^7^ purified leukocytes mouse 24 hours before back biopsy. To perform back biopsy wounds, mice were put under anesthesia using isoflurane. Mice were then given buprenorphine SR (sustained release) at 1 mg/kg via an IP injection before being moved to the surgical platform. Mice were measured from the nose cone edge to the base of tail and marked dorsally at the midpoint. Two biopsy sites were then marked at equal distances laterally from the midpoint, one on each side of the midline, after which mice were laid on their side and the loose skin was pulled taut. A 6-mm biopsy punch was then lined up with the surgical site marks and was pushed through the loose skin to create two holes of equal size. Mice were given 1 ml of 0.9% sterile saline via IP injection for additional hydration. The mouse was laid back on its stomach, the skin was stretched back to its normal position, and the animal was placed in a clean, warmed mouse box for recovery.

High-dimensional immunophenotyping of leukocyte invasion into back biopsy wounds was performed at 3 days post injury by the following methods. Adult (13–15-week-old) age-matched male nude mice were given saline or T-cell suspensions via the tail vein (adoptive transfer). Six-mm biopsy wounds were made, and the skin samples were harvested from euthanized mice by cutting the back skin in a square around both biopsy holes. An 8-mm biopsy was cut around the existing 3-day-old 6-mm biopsy wounds. The 2-mm ring of skin tissue was chopped with fine scissors and enzymatically digested in a gentleMACS C-tube using a skin digestion buffer (final enzyme concentrations:1 mg/mL collagenase type III [Worthington Biochemical Corporation], 0.33 mg/ml collagenase type 2 [Worthington Biochemical Corporation], 1.2 U/mL Collagenase/Dispase [Roche], 0.4 mg/ml of DNase I [Worthington Biochemical Corporation], and 2% FBS in HBSS with 0.9 mmol/L CaCl_2_). Samples were run on human skin program 1 on GentleMACS, and gentleMACS C-tubes were rocked for 90 minutes at 37°C. Samples were then run on the GentleMACS human skin program 1 before being filtered and flushed into a 50-ml conical tube with 40 ml of HBSS 5mM EDTA. The cells/debris were spun at 200 x g for 20 minutes (without brakes) before being resuspended in 250 µL of 2% FBS and HBSS, after which cells were counted.

Single-cell suspensions were treated with unlabeled Fc receptor-blocking antibody cocktail (CD16/CD32) in 50 µl of FACS buffer (HBSS + 5mM EDTA + 2% FCS). We then incubated 1×10^6^ cells in an antibody-staining cocktail against 20 cell-surface markers for 60 minutes at 4°C in 100 µl FACS Buffer. Cells were then washed in 1ml of FACS buffer and centrifuged at 400 x g for 7 minutes, 2 times. Cells were then resuspended and analyzed on a five-laser 30-parameter FACSymphony A5 cytometer (BD Biosciences, San Jose, USA) using DAPI exclusion for cell viability. For compensation of fluorescence spectral overlap, UltraComp eBeads (eBioscience, Inc.) were used following the manufacturer’s protocols. FCS 3.0 files generated by flow cytometry were initially processed using FlowJo Software (Tree Star, Ashland, OR) for automated compensation. Standard manual hierarchical gating was performed to remove debris, cell doublets, and (DAPI^+^) dead cells from analysis before gating on CD45^+^ leukocyte populations. In preparation for performing the 21-marker T-Distributed Stochastic Neighbor Embedding (tSNE) leukocyte-population analysis, the CD45^+^ population from each sample was down-sampled to 3,000 events to normalize cellular input between samples. Using FlowJo, a concatemer of all samples was performed. An unbiased tSNE plugin algorithm was then run, using defaults with 21 parameters (including DAPI channel), on the entire sample pool to obtain a multi-sample population reference map. Gating of each sample and experimental group was performed to generate tSNE maps for each condition. Differential cell clusters were gated, and 20-marker histogram plots were used to predict cluster identities.

Antibodies used for the 21 cell-surface-marker panel (purchased from Becton Dickinson (BD), New Jersey. USA or Biolegend, San Diego USA) are shown here: Anti-Mouse CD45-BUV395 (30-F11), Anti-Mouse B220_BUV496 (RA3-6B2), Anti-Mouse Ly-6G BUV563 (Clone 1A8), Anti-Mouse CD19 BUV661 (Clone 1D3), Anti-Mouse CD11b BUV737 (M1/70), Anti-Mouse CD4-BUV805 (RM4-5), Anti-Mouse NKp46_BV421 (29A1.4), Anti-Mouse CD8a BV510 (53-6.7), Anti-Mouse Ly6C_BV570 (HK1.4), Anti-Mouse Mrc1_BV650 (C068C2), Anti-Mouse MHCII-I-A/I-E -BV711 (M5/114.15.2), Anti-Mouse CD86-BV750 (GL1), Anti-Mouse CD11c BV786 (HL3), Anti-Mouse TCR delta -FITC (GL3), Anti-Mouse CD80 PE (16-10A1), Anti-Mouse CD115 PE-594 (ASF98), Anti-Mouse Cx3CR1 PercP5.5 (SA011F11), Anti-Mouse CD64 PE-Cy7 (X54-5/7.1), Anti-Mouse CD3e-APC (145-2C11), Anti-Mouse CCR2 A700 (475301), and Anti-Mouse F480 APC/Cy7 (BM8) .

## Acknowledgements

We acknowledge the JAX Center for Biometric Analysis, Pre-clinical Imaging division for the provision of instrumentation, training, and technical support in micro-CT imaging. Specifically, Doug Morris and Christine Wooley for project and instrument assistance. We acknowledge both the JAX Light Microscopy core, in particular Philipp Henrich for valuable support in 2 photon and confocal imaging along with MDI Biological Laboratory’s (MDIBL’s) Light Microscopy Facility (LMF) and Frederic Bonnet for widefield imaging assistance. We thank Will Schott, Krystal-Leigh Brown and Danielle Littlefield at JAX’s Flow Cytometry Service for their support and professional cell sorting assistance. We would like to thank Jax fabrication services, Jarek Trapszo in particular, for patent pending 3D printed mouse feet holder device (design and manufacture) used for advanced MicroCT imaging. We also would like to thank Stephen Sampson, Iain Drummond and Zhenying Jin for careful reading of the manuscript. We acknowledge the use of BioRender software for assistance in cartoon/figure design.

## Author contributions

Z.B. performed digit-tip surgeries, microscopy, micro-CT imaging and analysis, and adoptive transfer of purified cells, and assisted in data interpretation, manuscript preparation, and figure generation. R.R. performed digit-tip surgeries, histology, 2-photon and confocal microscopy and assisted with both cell purification and data analysis. L.H. helped with micro-CT imaging and analysis. N.R. provided infrastructure and manuscript editing. J.G. conceived the study, generated the hypothesis, designed experiments, performed high-dimensional flow cytometry and cell purification for adoptive transfers, analyzed data, and wrote/edited the manuscript.

## Data availability

The authors declare that all data supporting the findings of this study are available within the article and its supplementary information files, or from the corresponding author upon reasonable request.

## Supplementary Figures

**Figure S1.**
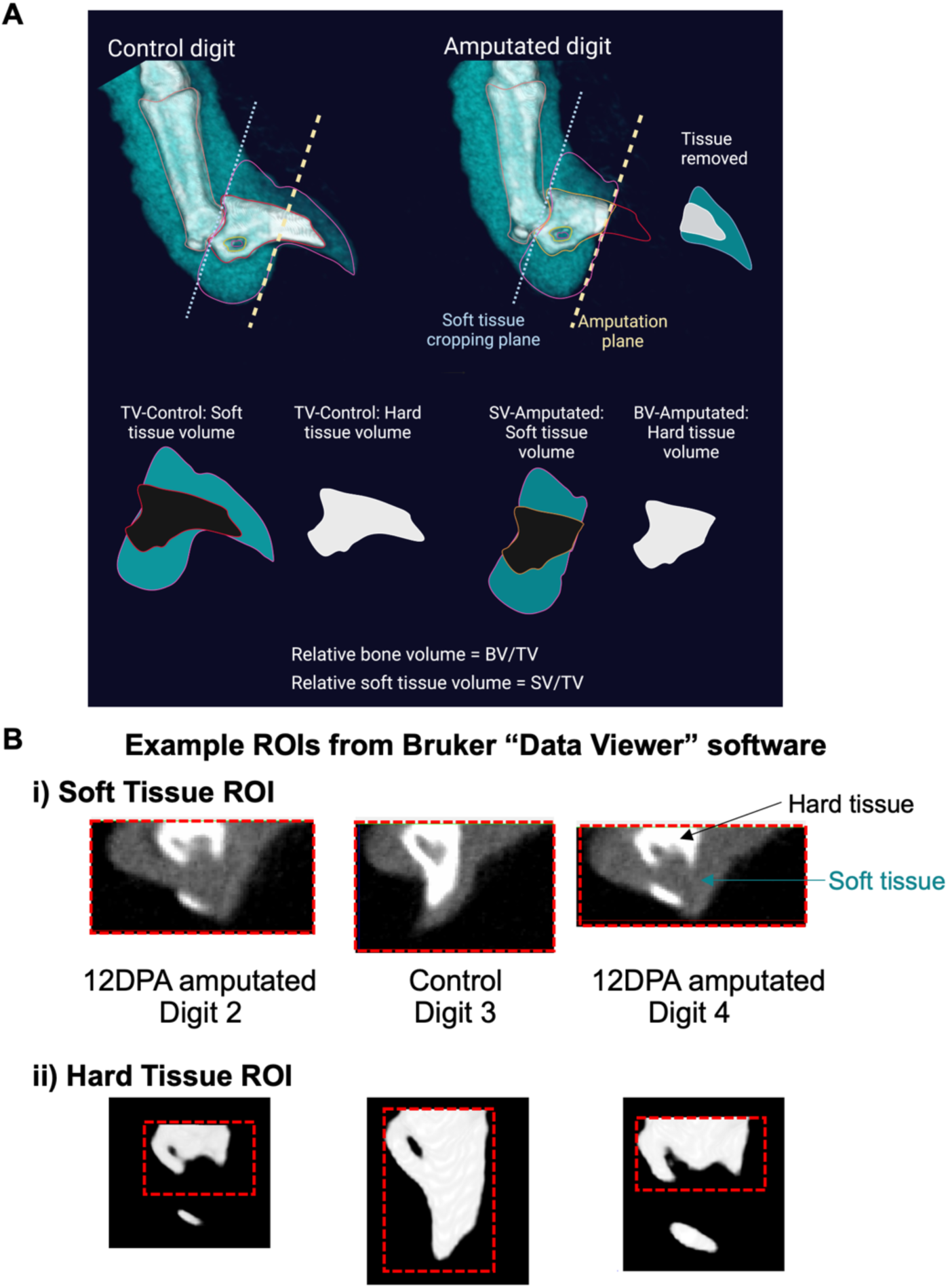
Two-tone quantification strategy. **A.** 2D picture of a typical 3D rendering of a mouse digit before and after P3-level amputation. Cartoon visualization of the soft tissues measured (teal) and the hard tissue (white) in both control and amputated digits. Relative bone volume calculated by dividing the regenerating digit (amputated) bone volume (BV) by the total volume (TV) of the unamputated control digit. Similarly, relative soft volume is calculated using SV/TV. Both relative values are converted to % of control. **B.** Examples of regions of interest (ROI) used to calculated 3D volumes in Bruker CTAn on microCT DICOM files shown in red dashed box. Note that in (i), the volume is calculated using the soft-tissue threshold (described in the Methods section) that excluded the volume of the hard tissue. Note that in (ii), only hard tissue contiguous with the main P3 bone element is included, with omission of any P3 fragment separated from the main P3 element during catabolism.

**Figure S2.**
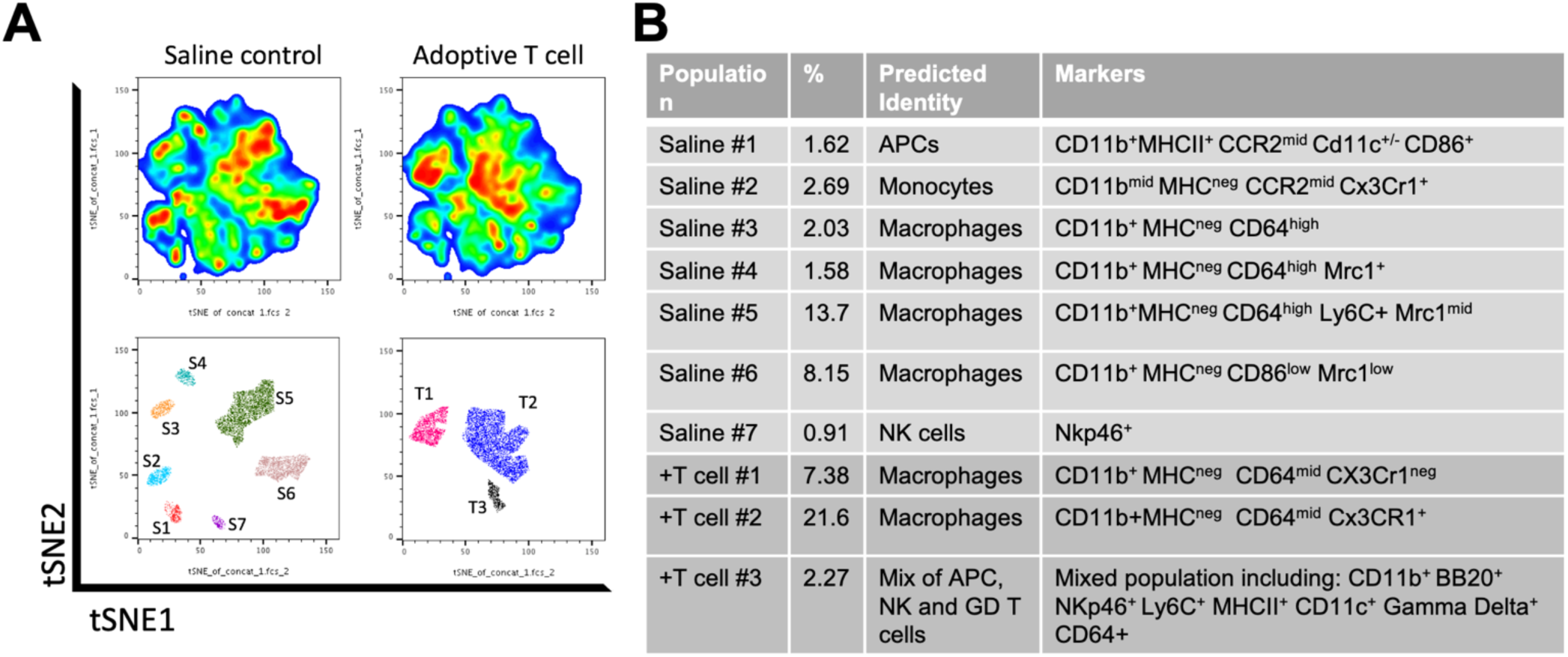
High-dimensional immune-phenotyping of skin biopsy wounds in T-cell-deficient nude (NU/J) mice receiving T-cell reconstitution shows dramatic changes in myeloid recruitment and differentiation. Dissociated skin biopsy samples 3-days post wounding were profiled using a 21-surface-marker flow cytometric panel to identify leukocyte changes after adoptive transfer of purified T cells. **A**. t-Distributed Stochastic Neighbor Embedding (tSNE) analysis was performed on CD45^+^ leukocytes normalized for input comparing saline injected and adoptive T cell treatments. **B.** Seven distinct major immune populations were identified in control (saline injected) animals (S1-S7), whereas only three major cell clusters could be identified in animals with T-cell reconstitution (T1-T3). The third column of the table in Fig. S2B shows the predicted identities of each cell cluster, using cell-surface markers.

**Figure S3.**
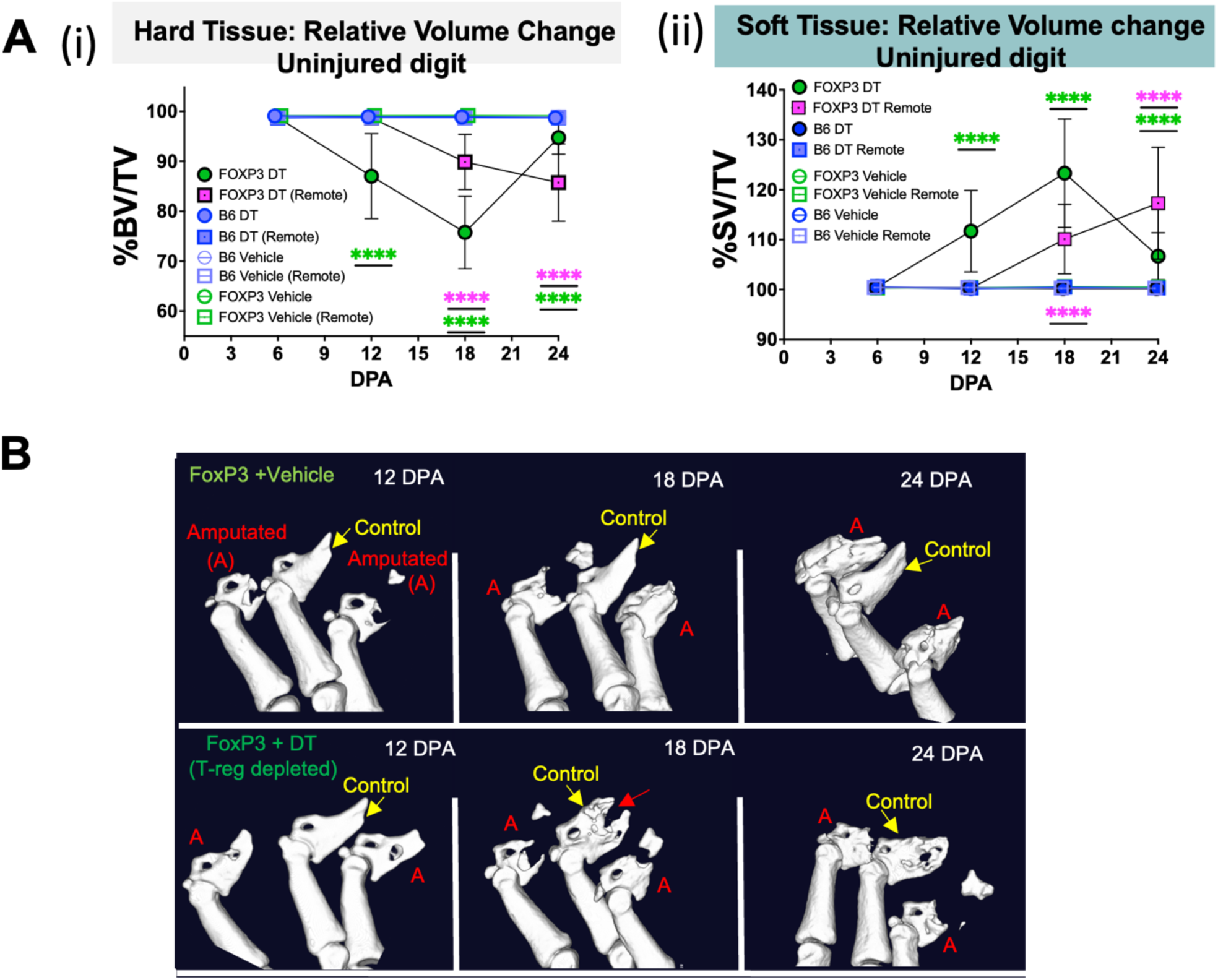
T-regs are necessary for preventing osteoclast-mediated autoimmunity and bone loss in the presence of other lymphoid cells. **A, B.** Sustained T-reg ablation results in erosion of control (uninjured) digits approximately 19 days from the first dose of DT. Erosion of bone is observed in both control (uninjured) digits on the same paw as amputated digits and to a lesser extent in distally “remote” uninjured front-paw digits. **A.** Changes in hard-tissue (**i**) and soft-tissue (**ii**) relative to uninjured control over time. Induced catabolism occurs in uninjured digits of both rear and distally remote uninjured front paws in T-reg-deficient animals. N = 5 mice with 4 amputated rear digits per animal = 20 measurements per condition, per timepoint. Data representative of at least 3 independent experiments. BV = bone volume, SV = soft volume, TV = total volume. P values calculated using one-way ANOVA with Turkey’s multiple comparison test. Statistical differences relative to B6 reference strain are indicated as ^ns^P > 0.05, *P ≤ 0.05, **P ≤ 0.01, ***P ≤ 0.001, ****P ≤ 0.0001 **B.** Representative single-color microCT images showing delayed bone catabolism in T-reg-deficient regenerating digits and osteoclastic attack of middle uninjured digit evident at 18 DPA (red arrow indicates site of osteoclastic attack). Amputated digits labelled with “A”. Unamputated middle control digits labelled in yellow with yellow arrow.

**Figure S4.**
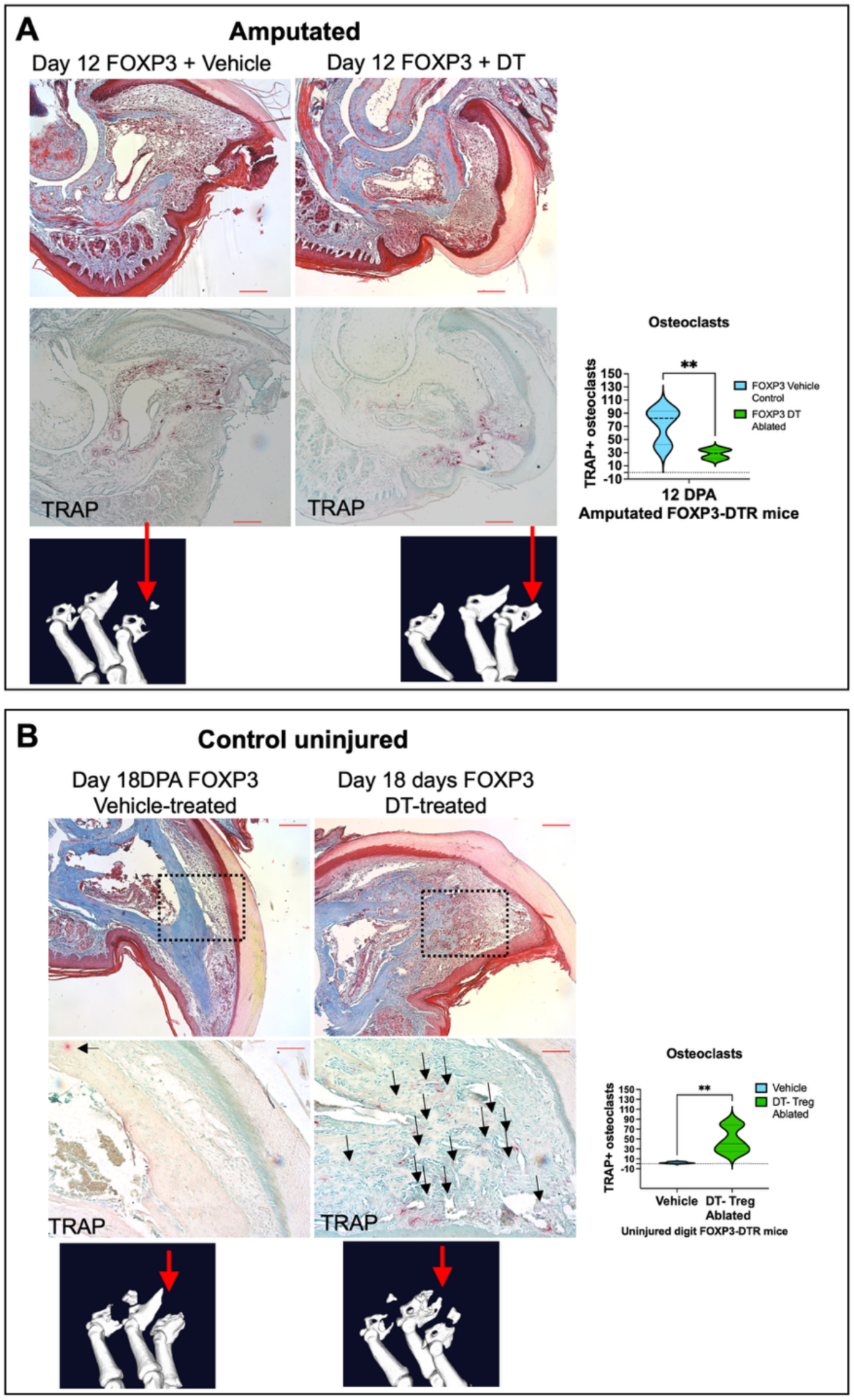
Diphtheria toxin (DT)-mediated depletion of genetically sensitized FoxP3^+^ T-regs delays the bone-catabolism phase of regeneration and induces dysregulation of bone homeostasis in uninjured digits. **(A)** Sustained DT-mediated ablation of T-regs in FoxP3-DTR mice results in disrupted digit-tip regeneration. Masson’s trichrome and TRAP staining of mature osteoclasts 12 DPA in amputated digits. Representative microCT rendering of bone indicates position of sections. (**B**) Sustained DT-mediated ablation of T-regs in FoxP3-DTR mice results in bone loss and increased numbers of TRAP^+^ osteoclasts in uninjured digits. Masson’s trichrome and TRAP staining of mature osteoclasts in non-amputated digits comparing animals treated with DT or vehicle. Uninjured digits harvested from animals with other regenerating digits at 18 DPA. P values obtained using one-way ANOVA with Turkey’s multiple comparison test. N = 3 biologically independent samples, averaging 3 sections per sample (taken around the midpoint of the digit). Means and standard deviations are shown. Statistical differences relative to vehicle control are indicated as ^ns^P > 0.05, *P ≤ 0.05, **P ≤ 0.01, ***P ≤ 0.001, ****P ≤ 0.0001. Scale bar = 100 μm

**Figure S5.**
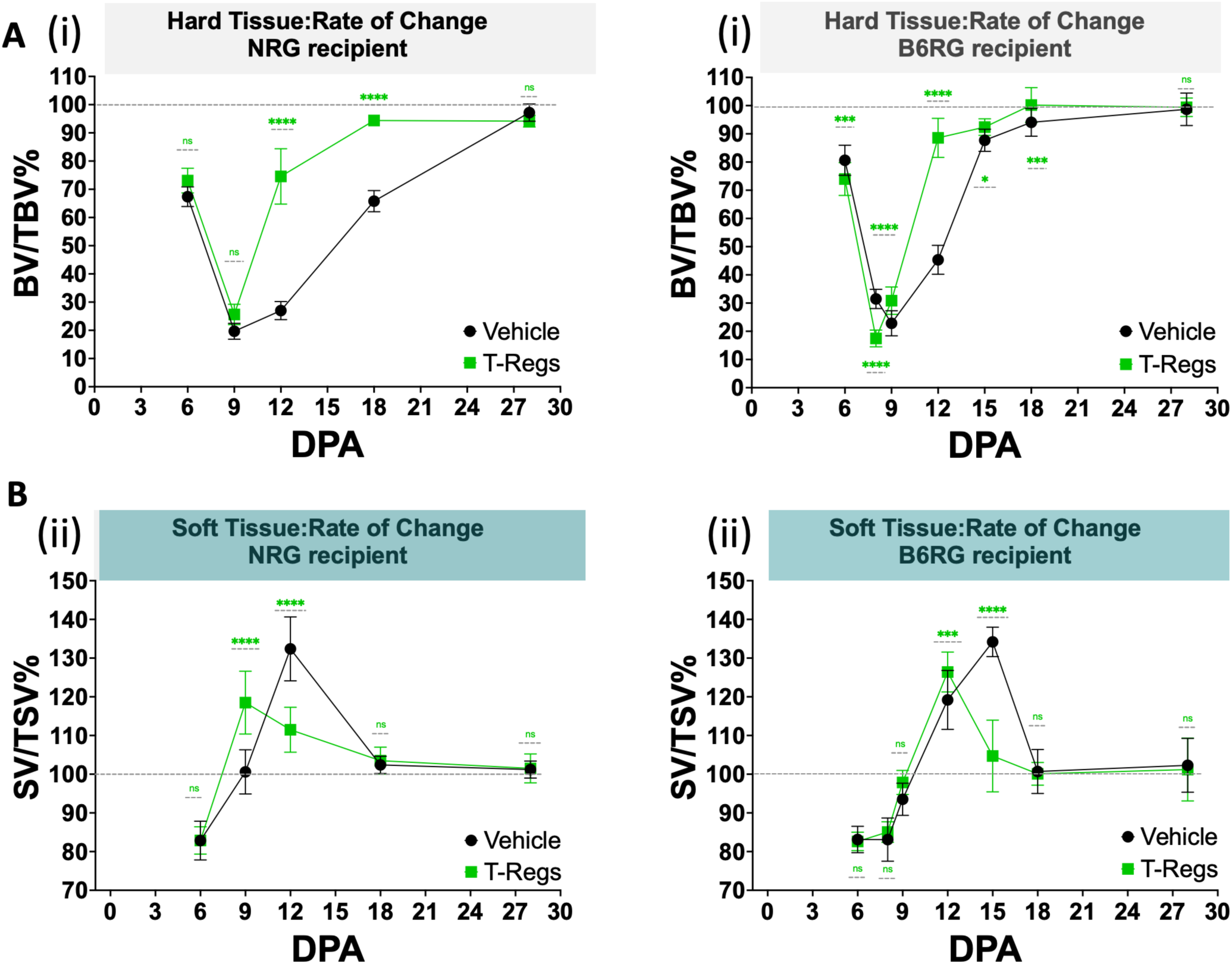
ACT of FoxP3^+^ T-regs in lymphoid-deficient mice enhances anabolic bone regeneration in two genetic backgrounds. FoxP3^+/YFP+^ T-regs were isolated via FACS and adoptively transferred into mice of the NRG and B6RG lymphoid-deficient strains one day prior to P3-level digit-tip amputations. **A.** Changes in hard tissue relative to uninjured control digits over time. N = 5 mice with 4 amputated digits per animal = 20 measurements per condition, per timepoint. Data representative of at least 2 independent experiments. BV= bone volume, SV = soft volume, TV = total volume. P values calculated using one-way ANOVA with Turkey’s multiple comparison test. Statistical differences indicated as ^ns^P > 0.05, *P ≤ 0.05, **P ≤ 0.01, ***P ≤ 0.001, ****P ≤ 0.0001. Means and standard deviations are shown. **B.** Changes in soft tissue relative to uninjured control digits over time. N = 5 mice with 4 amputated digits per animal = 20 measurements per condition, per timepoint. Data representative of at least 2 independent experiments. BV= bone volume, SV = soft volume, TV = total volume. P values calculated using one-way ANOVA with Turkey’s multiple comparison test. Statistical differences indicated as ^ns^P > 0.05, *P ≤ 0.05, **P ≤ 0.01, ***P ≤ 0.001, ****P ≤ 0.0001. Means and standard deviations are shown.

**Figure S6.**
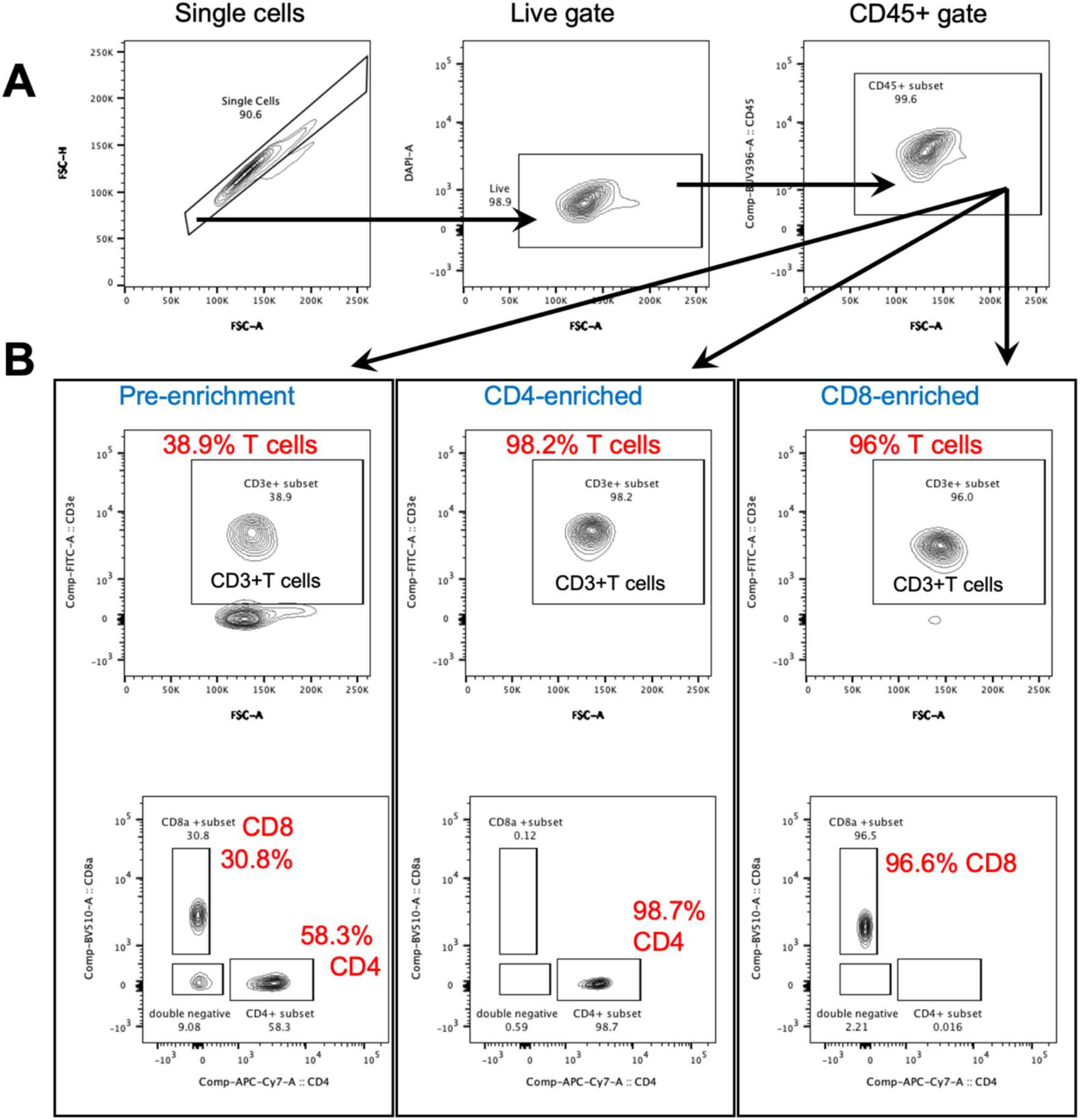
Example of BioLegend “MojoSort” magnetic-based enrichment of CD4^+^ and CD8^+^ T cells. **A.** Gating strategy for assessment of T-cell purity after enrichment of T cells from the mouse spleen. **B.** High levels of purity, i.e., over 98% for CD4^+^ T cells and 96% CD8^+^ T cells, were obtained from spleens, with an average of 28.9% total T-cell pre-enrichment.

**Figure S7.**
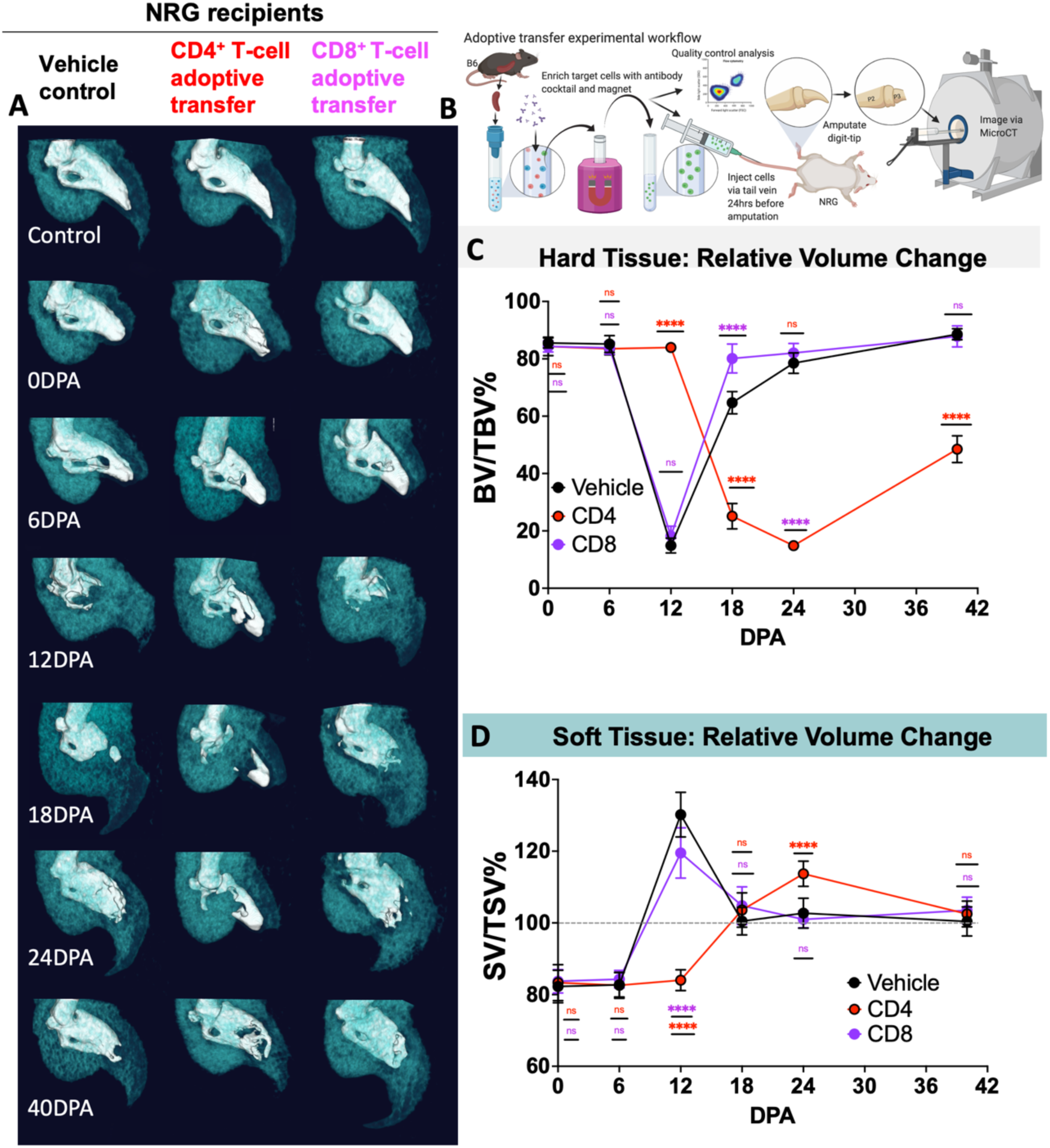
In NRG mice lacking lymphoid immunity, ACT of CD4^+^ T cells but not CD8^+^ T cells blocks digit-tip regeneration, affecting both hard- and soft-tissue growth. ACT of magnetically purified CD4^+^ or CD8^+^ T cells one day prior to P3-level amputation shows different outcomes in the NRG genetic background depending on the cell type transferred. **A.** Representative microCT image sequence of hard tissue (white) and soft-tissue overlay (teal) during digit-tip regeneration. **B.** Cartoon depicting experimental schema (generated in BioRender). **C**. Changes in hard tissue relative to uninjured control over time. **D**. Changes in soft tissue relative to uninjured control over time. N= 5 mice with 4 amputated digits per animal = 20 measurements per condition, per timepoint. Data representative of at least 2 independent experiments. BV = bone volume, SV = soft volume, TV = total volume. P values calculated using one-way ANOVA with Tukey’s multiple comparison test. Statistical differences relative to vehicle controls are indicated as ^ns^P > 0.05, *P ≤ 0.05, **P ≤ 0.01, ***P ≤ 0.001, ****P ≤ 0.0001. Means and standard deviations are shown.

**Figure S8.**
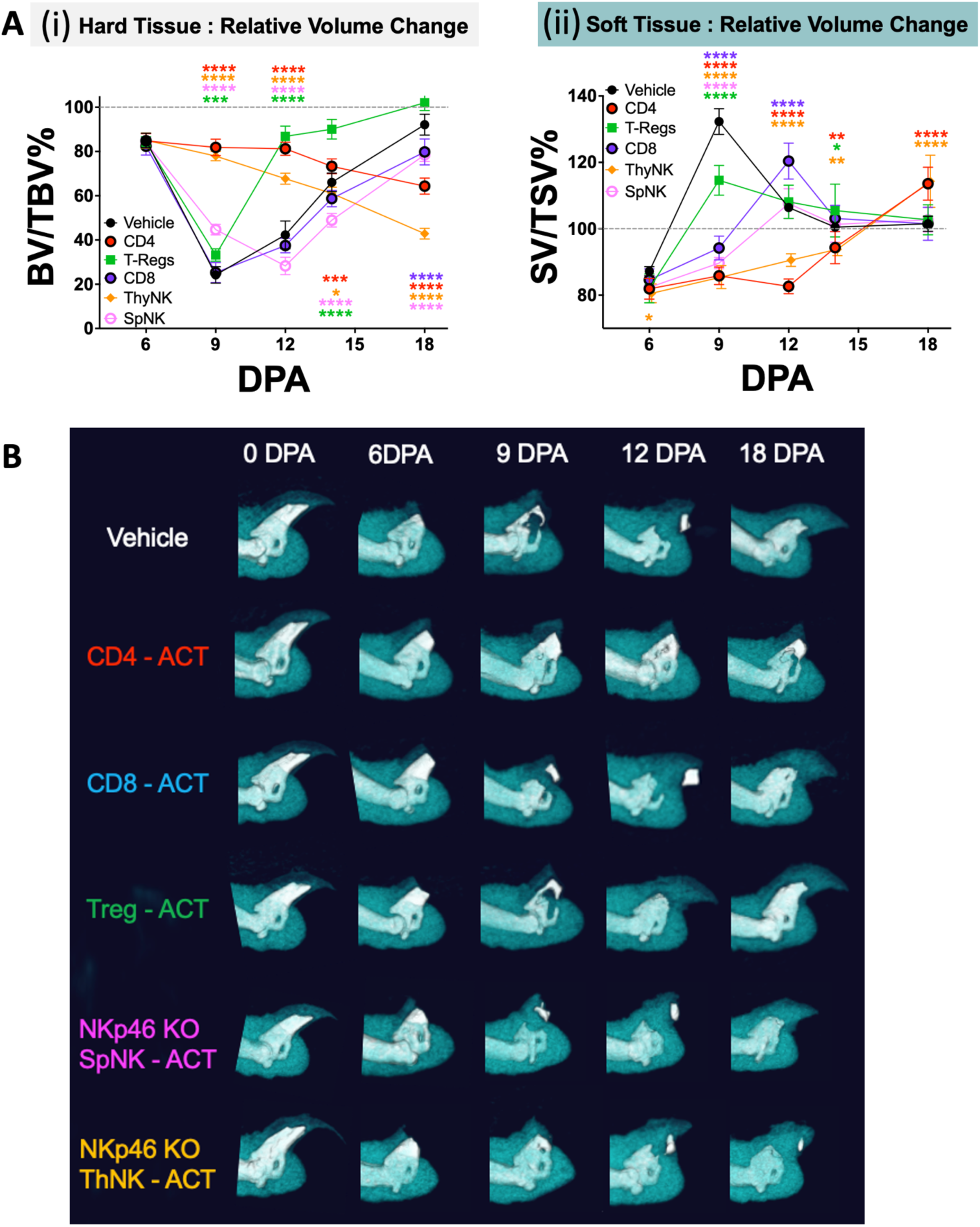
ACT of CD4+ T cells shows inhibitory effects on regeneration similar to those of thymic-derived NK cells. B6RG mice received magnetically (untouched purified CD4^+^ or CD8^+^ T cells or FACS-purified thymic-derived or splenic-derived NKp46 KI/KO GFP^+^ (Ncr1^gfp/gfp^) NK cells, or splenic T-regs (FoxP3^+YFP+^), or vehicle control, one day prior to P3-level digit-tip amputation. **A.** Changes in hard tissue (i) and soft tissue (ii) relative to uninjured control digits over time. N = 5 mice with 4 amputated digits per animal = 20 measurements per condition, per timepoint. Data representative of at least 2 independent experiments. BV = bone volume, SV = soft volume, TV = total volume. P values calculated using one-way ANOVA with Turkey’s multiple comparison test. Statistical differences indicated as ^ns^P > 0.05, *P ≤ 0.05, **P ≤ 0.01, ***P ≤ 0.001, ****P ≤ 0.0001. Means and standard deviations are shown. **B.** Representative two-tone microCT images of regeneration over time comparing treatments. Bone (hard tissue) shown in white, and soft tissue shown in teal.

**Figure S9.**
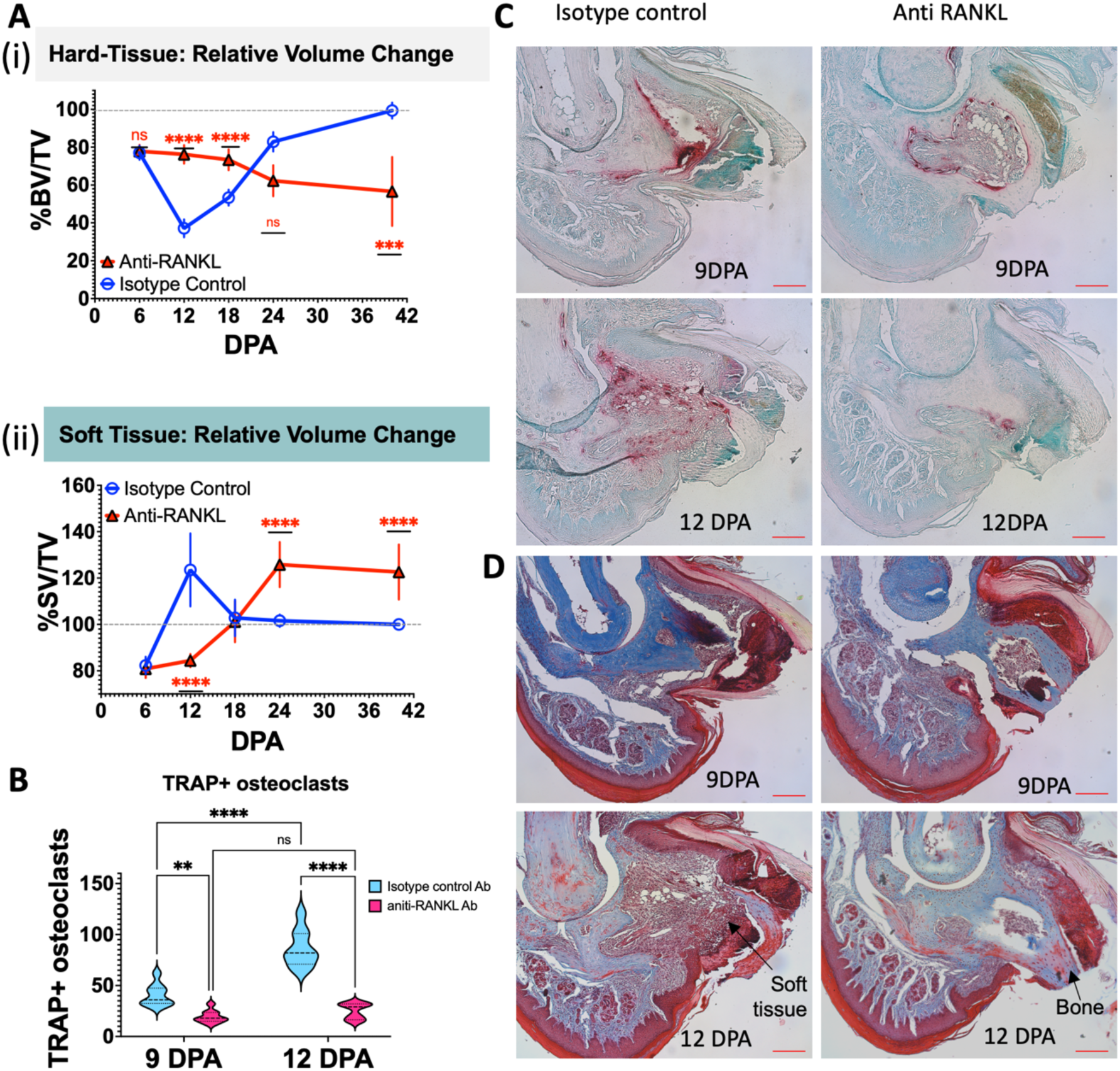
RANK ligand (RANKL) is necessary for the osteoclast-driven catabolic step that is required for digit-tip regeneration. Digits of fully immune-competent C57BL6/J (B6) control mice were amputated at the P3 position, and mice were injected IP with either *in vivo* grade anti-RANKL antibody or an isotype-matched control antibody at 6, 8, 10, and 12 DPA. Regeneration was monitored by microCT imaging. **A.** Volume of bone (hard tissue) (i) and soft tissue (ii) relative to uninjured control over time. N = 5 mice with 4 amputated digits per animal = 20 measurements per condition, per timepoint. Data representative of at least 2 independent experiments. BV = bone volume, SV = soft volume, TV = total volume. P values calculated using one-way ANOVA with Turkey’s multiple comparison test. **B**. Quantification of TRAP^+^ osteoclasts at 9 DPA and 12DPA with each treatment. **C**. TRAP staining of mature osteoclasts comparing anti-RANKL inhibited vs. control at 9 DPA and 12DPA with osteoclasts shown in red and nuclei counterstained in green. **D**. Masson’s trichrome staining comparing anti-RANKL-inhibited vs. control at 9 DPA and 12 DPA. N=3 biologically independent samples, averaging 3 sections per sample (taken around the midpoint of the digit). Data are represented as mean ± SD. P values less than 0.05 were considered significantly different from control (^ns^P > 0.05, *P ≤ 0.05, **P ≤ 0.01, ***P ≤ 0.001, ****P ≤ 0.0001). Means and standard deviations are shown. Scale bar = 100 μm.

**Figure S10.**
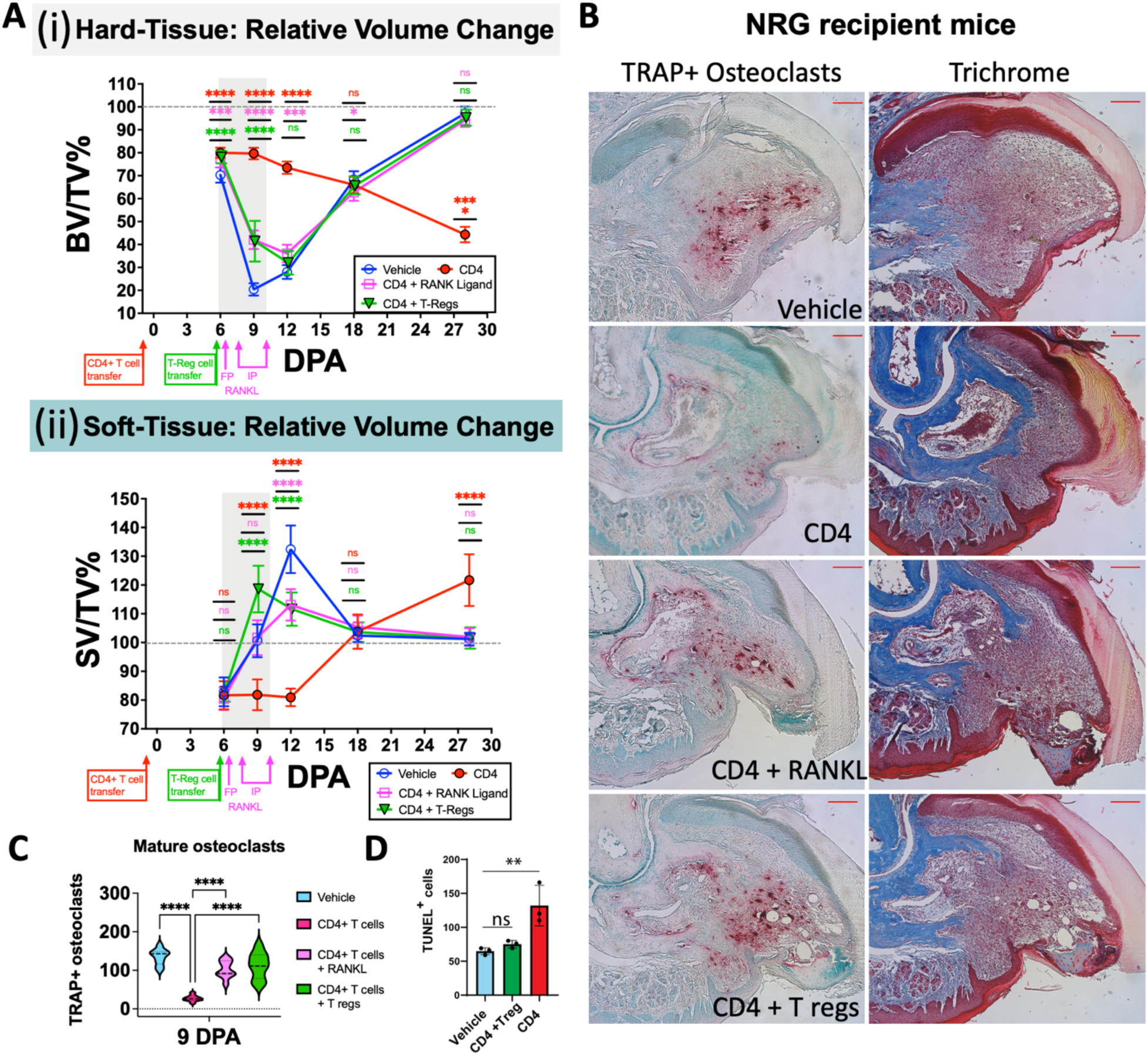
Asynchronous ACT of T-regs or delivery of recombinant RANKL in NRG mice rescues osteoclastogenesis and digit-tip regeneration in CD4^+^ T-cell-inhibited digits. Lymphoid-deficient NRG mice receiving purified CD4^+^ T cells one day prior to P3 amputation show strong inhibition of regeneration relative to vehicle controls. ACT of 1x 10^5^ FACS-purified FoxP3^+^ T-regs delivered via tail vein at 6 DPA (7 days post CD4^+^ T-cell delivery) completely rescued regeneration to normal control levels. Delivery of recombinant RANKL via footpad at 6 DPA and then via IP at 8 DPA and 10 DPA also rescued regeneration to normal levels in CD4^+^ T-cell-inhibited mice. **A.** Changes in hard-tissue (i) and soft-tissue (ii) relative to uninjured control over time. Timing of interventions indicated on X-axis, with RANKL treatment window shaded in grey. N = 5 mice with 4 amputated digits per animal = 20 measurements per condition, per timepoint. Data representative of at least 2 independent experiments. BV = bone volume, SV = soft volume, TV = total volume. P values calculated using one-way ANOVA with Turkey’s multiple comparison test. **B.** TRAP and Masson’s trichrome staining of osteoclasts at 9 DPA. Scale bar = 100 μm. **C.** Quantification of osteoclasts at 9 DPA. P values obtained using one-way ANOVA with Tukey’s multiple comparison test. **D.** Quantification of TUNEL+ (apoptotic) cells in each condition (Saline treated (vehicle) versus ACT of CD4^+^ T cells and asynchronous FoxP3^+^ T-regs and CD4^+^ T cell ACT) in NRG mice. N = 3 biologically independent samples, averaging 3 sections per sample (taken around the midpoint of the digit). Means and standard deviations are shown. P values calculated using one-way ANOVA with Turkey’s multiple comparison test. Scale bar = 100μm Statistical differences relative to B6 reference strain are indicated as ^ns^P > 0.05, *P ≤ 0.05, **P ≤ 0.01, ***P ≤ 0.001, ****P ≤ 0.0001.

